# Slow Outer Hair Cell Contractions Are Essential for Fast Electrical Responses to Intense Sound

**DOI:** 10.1101/2025.04.24.650391

**Authors:** Henrik Sahlin Pettersen, Anders Fridberger, Pierre Hakizimana

## Abstract

The cochlea protects itself from intense sound via slow mechanical contractions, but the real-time kinetics linking this process to the stimulus-evoked electrical potentials have remained unresolved. Here, using high-speed confocal imaging with AI-driven analysis, we synchronously measured organ of Corti mechanics and stimulus-evoked potentials in the living isolated guinea pig cochlea. We discovered an inverse kinetic relationship: the outer hair cell’s (OHC) slow, somatic contraction is essential for generating a fast electrical response. Pharmacologically blocking the OHC motor protein prestin inverted this dynamic; the OHC’s mechanical contraction became faster, while the normally rapid electrical potential became nearly ten times slower. These findings indicate that slow OHC motility is not merely a byproduct of overstimulation but a control mechanism. It functions to regulate and sharpen the kinetics of the stimulus-evoked potential, providing a cellular-level explanation for how the hearing organ protects itself while maintaining temporal fidelity during intense sound exposure.

## Introduction

Mammalian hearing achieves remarkable sensitivity and frequency selectivity through a process of active mechanical amplification within the cochlea [1]. Central to this process are the outer hair cells (OHCs), which physically amplify sound vibrations through somatic electromotility, a voltage-driven change in cell length powered by the motor protein prestin [2–6]. While OHCs are the engine of amplification, their function is intimately tied to the surrounding structures. Recent work has revealed that the cochlear partition is a more tightly integrated mechanical unit than classically described, with even the inner hair cell (IHC) stereocilia being physically embedded in the overlying tectorial membrane (TM) via novel, Ca^2+^-rich filamentous structures [7].

When this integrated system is challenged by intense sound, it engages protective mechanisms that lead to temporary shifts in hearing sensitivity. Foundational studies showed that acoustic overstimulation triggers slow, reversible contractions of the organ of Corti [8], linked to a rise in OHC intracellular Ca^2+^ [8, 9], and a concurrent depletion of the TM’s extracellular Ca^2+^ reservoir [10]. This acoustic stress is also encoded in the cochlea’s DC electrical response, or summating potential (SP). We recently showed that the SP serves as a direct biomarker of cochlear health, switching its polarity from positive in a healthy state to negative following noise-induced injury, which signals a shift in the hair cells’ operating point [11].

Together, these studies reveal that intense sound induces a complex response involving organ-level mechanics, ionic shifts, and a distinct electrical signature. However, the kinetic relationship between the OHC’s own motor activity and the resulting stimulus-evoked potential during acoustic stress has remained unresolved. Resolving whether OHC motility merely follows the electrical potential or regulates it—a crucial distinction for understanding cochlear protection—requires synchronous measurement of both phenomena, a longstanding technical challenge. In this study, we overcame this barrier using AI-enhanced high-speed imaging with simultaneous electrophysiology in the living isolated guinea pig cochlea to dissect this interplay and describe a novel principle of cochlear control.

## Results

To investigate the relationship between outer hair cell (OHC) mechanical responses and the kinetics of stimulus-evoked potentials during acoustic stress, we developed an integrated experimental platform for synchronous mechanical and electrical recording in the nearly intact guinea pig cochlea (Fig. 1). Using a stepped acoustic stimulation protocol that included baseline, moderate, and intense sound phases, we simultaneously captured high-speed confocal image series to quantify OHC mechanical changes while recording the corresponding stimulus-evoked potentials. The silent (0 dB SPL) baseline and recovery periods were included to monitor the mechanical stability of the preparation; as demonstrated in Supplementary Fig. S1, the OHC area change returned to a stable baseline, confirming the preparation’s viability. This approach allowed us to directly correlate the kinetics of OHC somatic contractions with the underlying electrical signals on a millisecond-to-second timescale.

**Figure 1:**
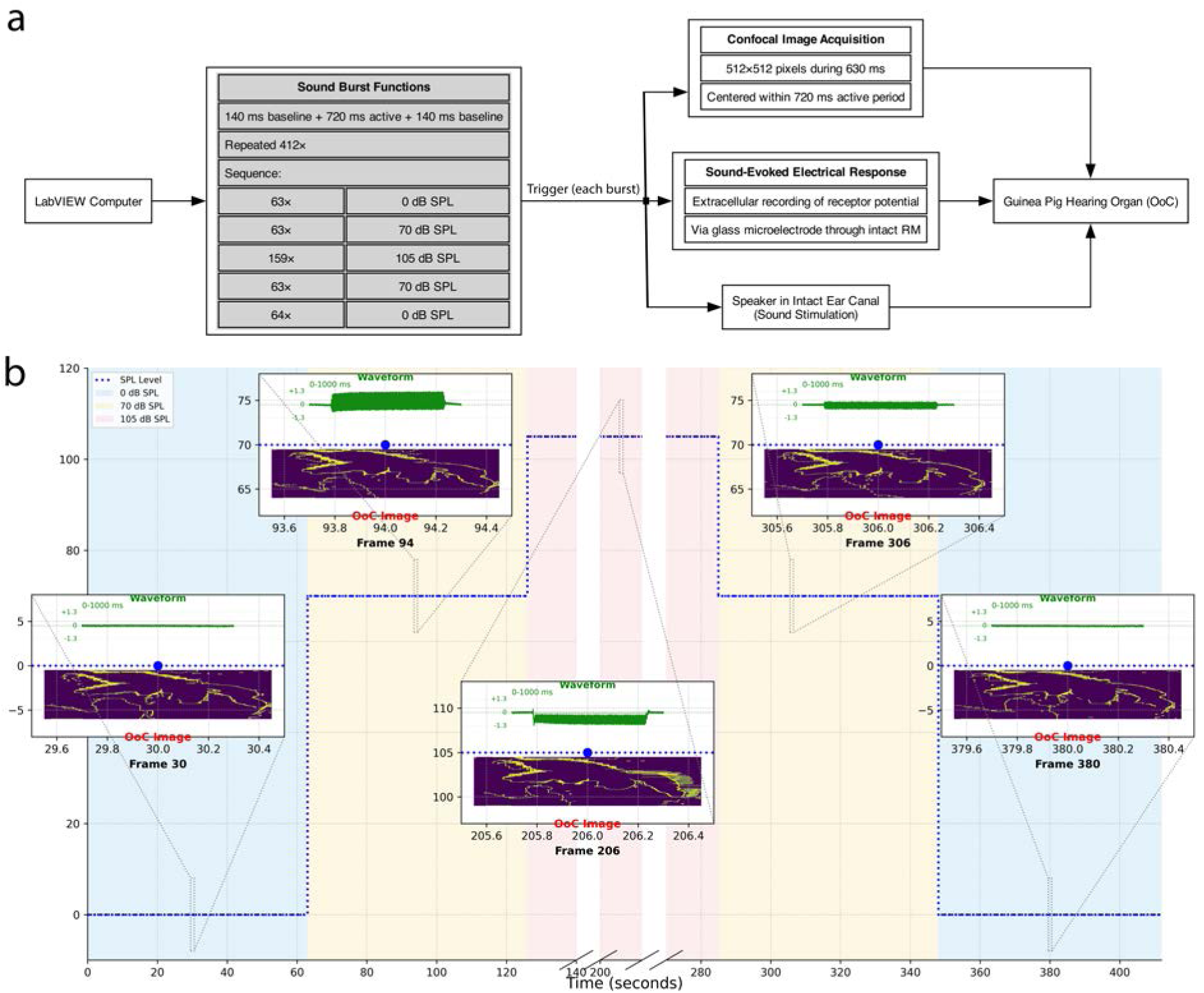
Experimental design for synchronous mechano-electrical recording. **a,** Schematic of the experimental platform. A LabVIEW computer orchestrates the synchronized acquisition of confocal images and sound-evoked potentials from the guinea pig hearing organ. **b,** Experimental timeline consisting of 412 one-second tone bursts at 160 Hz (total duration 412 s), showing the stepped acoustic stimulation protocol. The protocol comprises five phases defined by sound pressure level: initial baseline (0 dB SPL), moderate sound (70 dB SPL), intense sound (105 dB SPL), and subsequent recovery periods. Insets provide representative examples from the control dataset, showing a characteristic electrical waveform (in mV) and Organ of Corti (OoC) image for different phases of the protocol, with the latter displayed as a simplified contour for visual clarity.

### Prestin-dependent motility governs the dynamics of the SP response

In response to intense 160 Hz sound (105 dB SPL), the cochlea generated a robust, negative DC summating potential (SP)—a known signature of cochlear stress [11]. The SP exhibited a biphasic response, characterized by a rapid-onset component followed by a slowly developing, sustained decay over tens of seconds (Fig. 2). A quantitative summary of these electrical responses and the statistical outcomes of pharmacological manipulations is presented in Supplementary Fig. S2.

**Figure 2:**
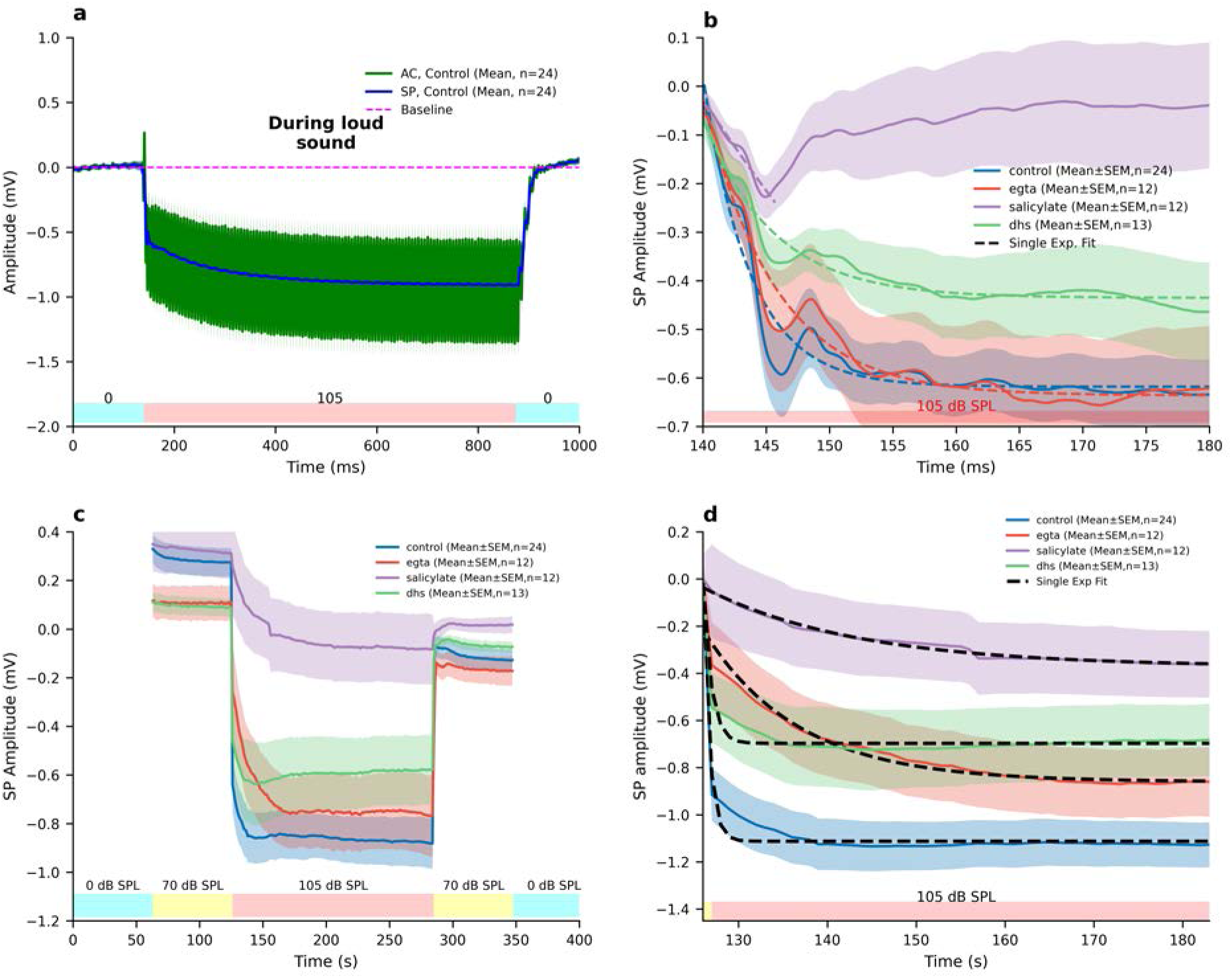
Prestin-dependent motility regulates the biphasic kinetics of the summating potential. **a,** The averaged electrical recording from control cochleae (*n* = 24) during intense sound (105 dB SPL) shows the alternating current (AC, green) and direct current (DC) summating potential (SP, blue) components. Dashed magenta line indicates the pre-stimulus baseline. **b,** The rapid-onset component of the SP during the initial 40 ms of stimulation. Exponential fitting of the mean traces revealed a time constant (*τ*) of 3.7 ms in control conditions. Blocking prestin-dependent motility with salicylate (purple, *n* = 12) slowed these kinetics nearly tenfold to *τ* = 34.0 ms. **c,** Full time course of the SP showing the development of the sustained component of the response across multiple sound pressure levels. **d,** The sustained, slowly developing SP component during intense stimulation. The decay was fastest in control (*τ* = 0.7 s) and was significantly slower with salicylate (*τ* = 17.2 s) and EGTA (*τ* = 10.6 s). All data are presented as mean ± s.e.m. (shaded bands).

To dissect the underlying mechanisms, we first analyzed the kinetics of the rapid-onset component by fitting a single exponential model to the mean traces. In control experiments (*n* = 24), this initial phase had a time constant of *τ* = 3.7 ms. Blocking OHC somatic contraction with salicylate (*n* = 12) dramatically slowed these kinetics, increasing the time constant nearly tenfold to *τ* = 34.0 ms (Supplementary Fig. S2c). In contrast, neither the chelation of extracellular Ca^2+^ with EGTA (*τ* = 5.6 ms) nor a partial block of mechanoelectrical transducer (MET) channels with DHS (*τ* = 5.5 ms) produced kinetics statistically different from the control condition (Supplementary Fig. S2c).

A closer inspection of the traces (Fig. 2b) reveals a brief oscillatory feature between approximately 147 and 150 ms in the control, EGTA, and DHS conditions. This feature is notably absent in the salicylate-treated cochleae, which instead exhibit a smoother, monotonic decay during this time window. While the exponential model is used to characterize the overall decay trend, it is important to acknowledge that this brief, systematic oscillation will necessarily influence the precise parameters of the fit. Nonetheless, the disappearance of this rapid oscillation upon blockade of the OHC motor strongly suggests that it is a direct electrical signature of prestin-dependent mechanical activity at the onset of intense stimulation. Beyond the rapid-onset component, the kinetics of the sustained SP decay were also strongly affected by pharmacological manipulation.

The sustained decay was most rapid in the control condition (*τ* = 0.7 s), with similar kinetics observed following a partial block of MET channels with DHS (*τ* = 0.9 s). In contrast, the decay became markedly slower upon chelation of Ca^2+^ with EGTA (*τ* = 10.6 s) and was slowest when prestin-dependent motility was blocked with salicylate (*τ* = 17.2 s) (Supplementary Fig. S2d). Statistical analysis of the final response amplitudes confirmed significant differences between control and both salicylate (*p* = 0.0002) and DHS (*p* = 0.0156), but not EGTA (*p* = 0.1241) (Supplementary Fig. S2b). These results demonstrate that while MET channels and calcium levels influence the SP, prestin-dependent motility is the principal regulator of the kinetics of both the rapid-onset and sustained components of the electrical response to intense sound.

To directly test how these altered electrical kinetics relate to the underlying cell mechanics, we compared the normalized electrical and mechanical response waveforms (Supplementary Fig. S3). In control conditions, the two signals were exceptionally well-coupled, with the slow OHC contraction tracking the development of the SP with a Pearson correlation coefficient of *r* = 0.926 (*p <* 0.001). Crucially, a strong temporal coupling was preserved even when prestin was blocked with salicylate (*r* = 0.710, *p <* 0.001), indicating that the two phenomena remain inextricably linked. This establishes that the mechanical response is inextricably linked to the electrical potential, setting the stage for investigating the mechanical basis of the profound kinetic shift observed in the stimulus-evoked potential.

### Slow OHC somatic contraction is accelerated when prestin is blocked

Having established a tight temporal link between the stimulus-evoked potential and cell mechanics, we next investigated the mechanical basis for the observed electrical kinetics by quantifying the cross-sectional area of OHCs during acoustic stimulation (Fig. 3). To generate the mechanical response waveforms, morphological parameters like OHC area were extracted from each frame of the AI-segmented videos. For each experiment, the resulting time-series was baseline-corrected by subtracting its mean value during the initial silent period before being averaged with other experiments. The full time-course of the organ of Corti’s mechanical response is visualized in the denoised confocal recordings (Supplementary Movie 1), and the output of our AI-driven tracking is shown in the corresponding segmentation video (Supplementary Movie 2).

**Figure 3:**
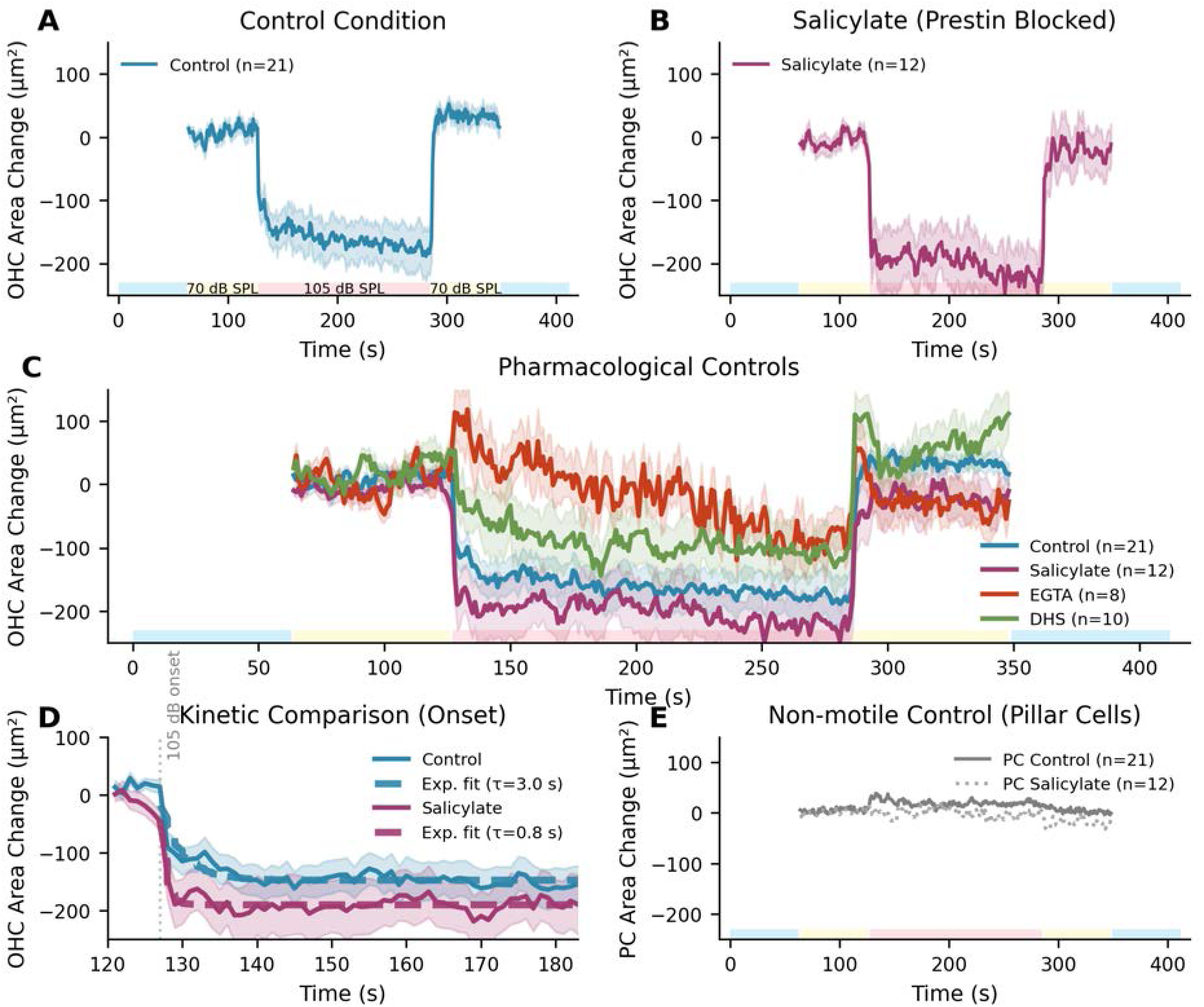
OHC somatic contraction kinetics are inversely regulated by prestin-dependent motility. **a**, In control conditions (*n* = 21), outer hair cells (OHCs) exhibited a slow, sustained contraction during 105 dB SPL stimulation, reaching a peak area reduction of 166 µm^2^. **b**, Blocking prestin with salicylate (*n* = 12) resulted in a faster and larger contraction (peak area reduction of 205 µm^2^). **c**, The contraction was significantly reduced by the Ca^2+^ chelator EGTA (*n* = 8, 80% inhibition) and partially reduced by the MET channel blocker DHS (*n* = 10, 42% inhibition). **d**, A direct comparison of the contraction onset reveals a 3.6-fold acceleration in kinetics for salicylate-treated OHCs. Single exponential fits show the time constant (*τ*) decreased from 3.0 s in control to 0.84 s with salicylate (*p <* 0.001). **e**, The non-motile Pillar Cells (PCs) showed negligible area changes (*<*25 µm^2^), confirming the response is specific to OHCs. Data are presented as mean ± s.e.m. Shaded bars on the x-axis indicate stimulus periods. All statistical comparisons are to the control condition.

To ensure the fidelity of this mechanical analysis, all image series underwent a stringent quality control protocol after acquisition. This process led to the exclusion of a small number of time series (3 of 24 in control, 4 of 12 in EGTA, 3 of 13 in DHS, and 0 of 12 in salicylate) due to factors such as low image contrast or motion artifacts, which can compromise the accuracy of automated segmentation. This necessary quality control step accounts for the difference in sample size (*n*) between the electrical and mechanical datasets.

The resulting analysis shows that area change arises from anisotropic changes in both cell length and width (Supplementary Fig. S4). In control conditions, exposure to intense sound (105 dB SPL) induced a slow somatic contraction in OHCs, characterized by a time constant of *τ* = 3.0 s and reaching a peak area reduction of 166 µm^2^.

Blocking prestin-dependent motility with salicylate—the same manipulation that slowed the electrical response (Fig. 2)—had the opposite effect on OHC mechanics. The OHC contraction became significantly faster, with its onset time constant decreasing 3.6-fold from *τ* = 3.0 s to *τ* = 0.84 s (*p <* 0.001, Fig. 3d; Supplementary Fig. S5b). The peak contraction magnitude also appeared larger with salicylate (205 µm^2^ vs. 166 µm^2^), though this difference was not statistically significant (*p* = 0.082, Supplementary Fig. S5a). This finding reveals an inverse kinetic relationship: disabling the OHC’s motor function accelerates its mechanical contraction but slows the stimulus-evoked potential. This suggests that the OHC’s normally slow contraction is a process that is required for a fast electrical potential. Following stimulation, control cells exhibited a slight swelling during recovery, whereas salicylate-treated cells showed residual shrinkage (Supplementary Fig. S1).

The specificity of this mechanism was confirmed with pharmacological and structural controls. The contraction was dramatically attenuated by the Ca^2+^ chelator EGTA (80% inhibition, *p <* 0.001) and partially reduced by the MET channel blocker DHS (42% inhibition, *p <* 0.001) (Supplementary Fig. S5). Furthermore, neighboring non-motile pillar cells showed minimal area changes (*<* 25 µm^2^, Fig. 3e), and a broader survey confirmed that the large mechanical response was unique among cochlear cell types (Supplementary Fig. S6).

### OHC somatic contraction and bulk displacement are distinct mechanical processes

To distinguish the OHC’s intrinsic shape change from the passive bulk movement of the entire cochlear partition, we analyzed two separate mechanical metrics: the change in the OHC’s cross-sectional area (an indicator of somatic contraction, Fig. 3) and the change in the cell’s overall position (axial displacement, Fig. 4). In response to intense sound, we observed a coordinated downward displacement across the organ of Corti, with OHCs exhibiting the largest peak amplitude (≈ −7.2 µm in control) (Supplementary Fig. S7, Supplementary Fig. S8). This movement was predominantly in the axial direction, which was consistently 4-to-7-fold greater than the radial component across all conditions (Supplementary Fig. S9).

**Figure 4:**
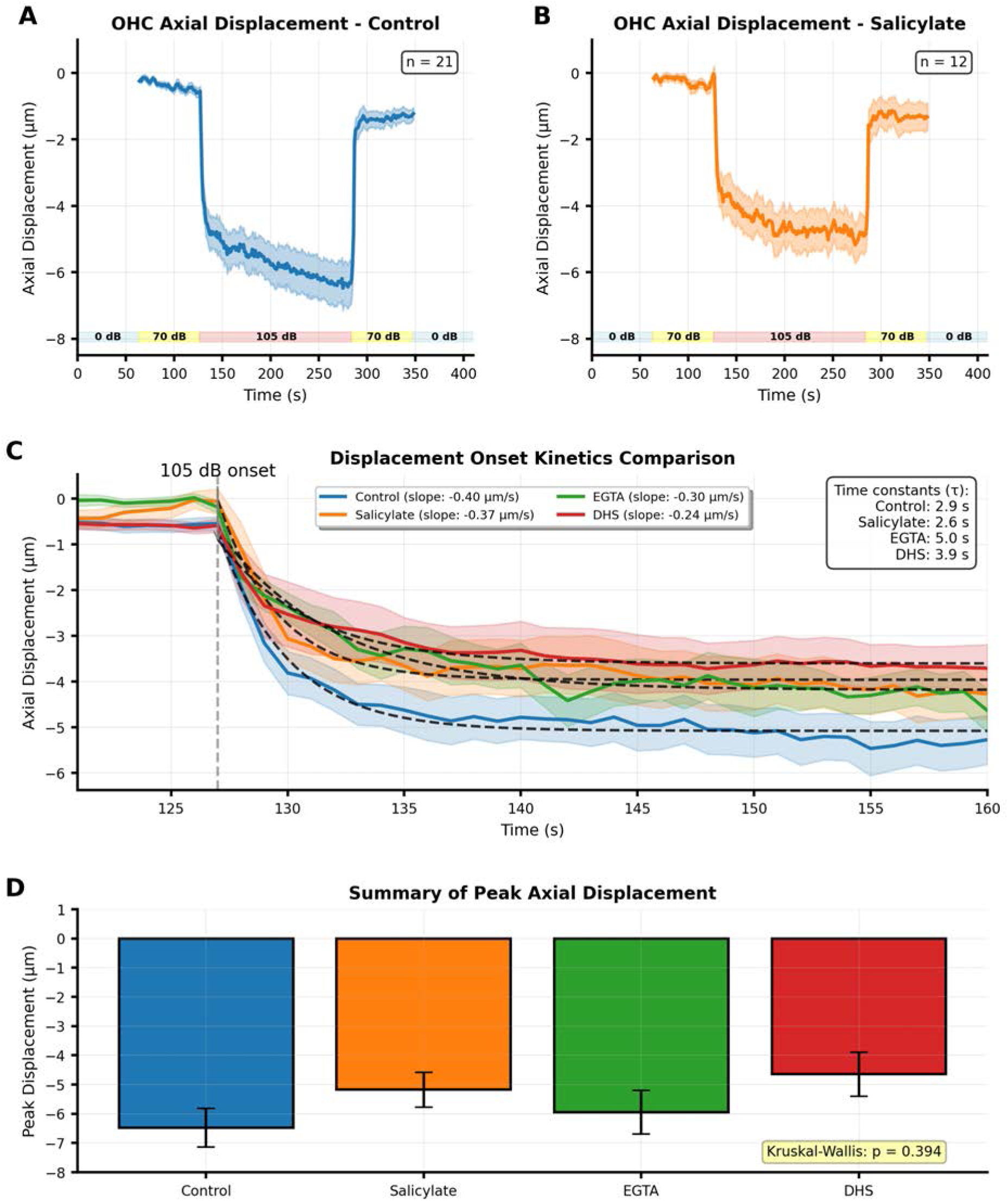
OHC displacement dynamics are distinct from somatic contraction kinetics. **a**, Time course of OHC axial displacement in control conditions (*n* = 21) showing slow, downward movement during the 105 dB SPL stimulus. **b**, OHC axial displacement following salicylate treatment (*n* = 12). **c**, Direct comparison of displacement onset kinetics. The onset slopes are remarkably similar, particularly between control (−0.40 µm*/*s) and salicylate (−0.37 µm*/*s), demonstrating that bulk displacement kinetics are not regulated by prestin in the same manner as somatic contraction. **d**, Summary of peak axial displacement magnitudes. Despite apparent differences in the mean, a Kruskal-Wallis test found no statistically significant difference across conditions (*p* = 0.394). Data in (a, b) are mean ± s.e.m.

Crucially, the kinetics of this bulk displacement did not show the same inverse relationship with prestin function that was seen with somatic contraction. The onset slope of OHC displacement was nearly identical between control (−0.40 µm*/*s) and salicylate-treated (−0.37 µm*/*s) conditions (Fig. 4c). Furthermore, while pharmacological treatments appeared to reduce the mean peak displacement, these changes were not statistically significant for OHCs (Kruskal-Wallis test, *p* = 0.394, Fig. 4d; Supplementary Fig. S8).

These results demonstrate a clear dissociation between two distinct mechanical processes. While somatic contraction is a finely tuned process whose kinetics are regulated by prestin, the bulk displacement of the cell appears to be a passive consequence of overall organ of Corti motion, whose kinetics and amplitude are not subject to the same control. This reinforces that the kinetic sharpening of the stimulus-evoked potential is specifically linked to the intrinsic shape change of the OHC.

### Cycle-by-cycle analysis validates OHC area as a robust metric for motility

Finally, to confirm that our area-change metric was a valid proxy for sound-evoked OHC motility, we performed high-resolution, time-resolved imaging [12, 13] of OHCs during moderate sound stimulation (70 dB SPL). High-magnification imaging provided clear anatomical context for the targeted OHCs (Fig. 5a-f). Using a pixel-clock-synchronized acquisition system [12, 13] and advanced denoising [14], we resolved nanoscale OHC displacements and area changes on a cycle-by-cycle basis (Fig. 5g, h, i). For details on the imaging and processing methodology, see Supplementary Fig. S10. OHCs exhibited consistent, cyclic area changes of approximately 2.7% relative to the steady-state area, an amplitude consistent with established estimates of OHC length changes [2, 3, 6, 15]. This high-temporal-resolution result confirms that our AI-powered area analysis is a sensitive and valid metric for quantifying OHC motility.

**Figure 5:**
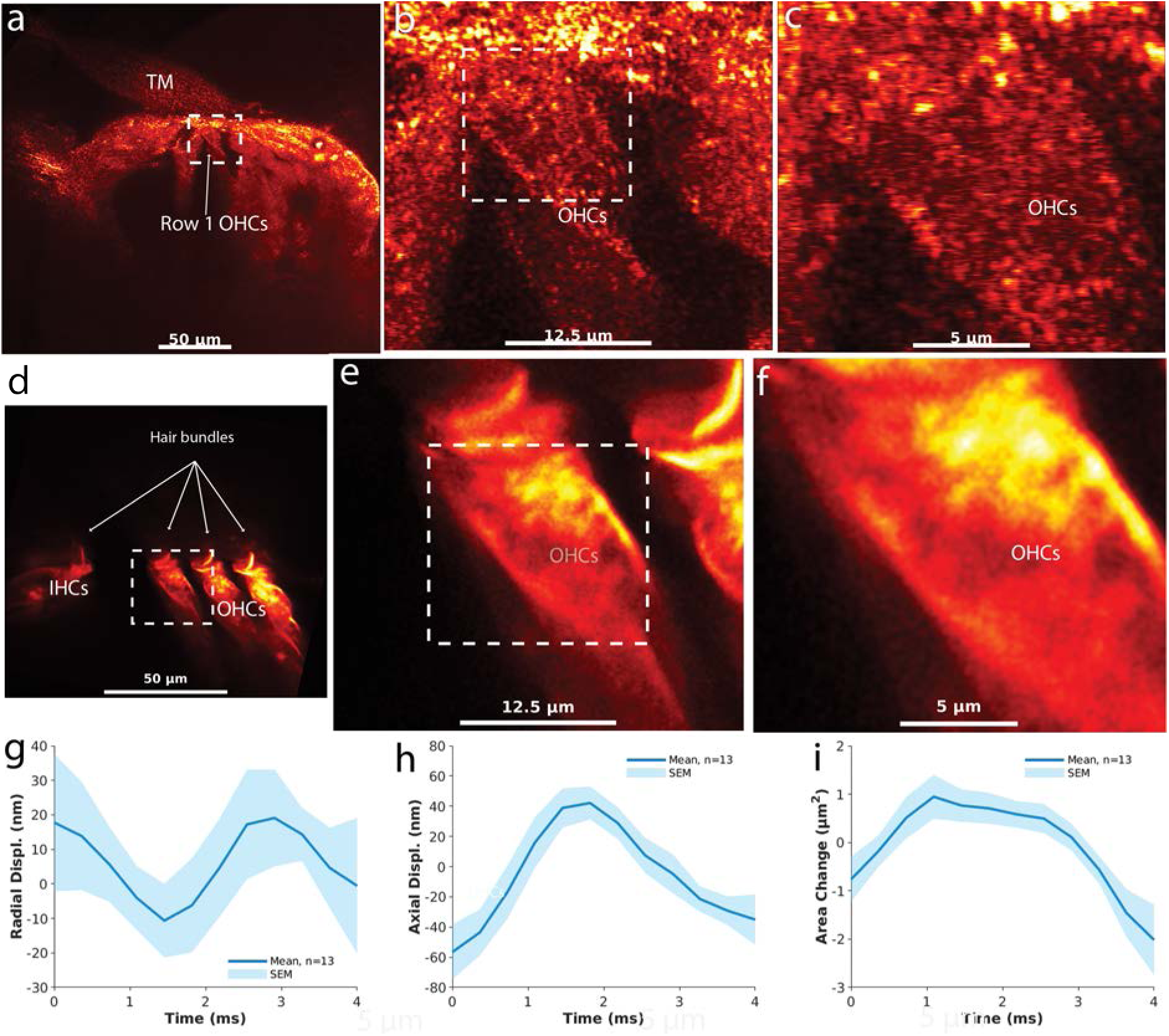
Cycle-by-cycle analysis validates OHC area change as a robust metric for motility. (**a-f**) High-magnification images providing anatomical context of the outer hair cells (OHCs) within the organ of Corti from which high-speed recordings were made. (**g-i**) High-speed imaging resolved OHC mechanical changes during one cycle of a 70 dB SPL stimulus (*n* = 13). The analysis captured cyclic changes in radial displacement (**g**), axial displacement (**h**), and cross-sectional area (**i**). The clear, periodic signal in the area measurement validates its use as a robust proxy for OHC motility in the main findings.

## Discussion

In this study, we have shown that a slow mechanical action from the outer hair cell (OHC) is necessary to produce a fast electrical response to intense sound. We propose a model where this slow, prestin-dependent contraction acts as a protective control system. **Conceptually, in the absence of slow, active OHC resistance, a more compliant partition could permit larger passive motions that shift the MET operating point and slow the receptor potential; this ‘mechanical damping’ framing clarifies how a slow process preserves a fast onset.** When we pharmacologically disabled this function with salicylate, the OHC’s underlying passive mechanical properties were unmasked, resulting in a faster somatic contraction at the cost of electrical fidelity. This inverse relationship indicates that the OHC’s slow contraction is a regulated process for signal conditioning and protection, not merely a byproduct of overstimulation.

It is important to acknowledge that salicylate has a dual pharmacological action on the OHC. In addition to blocking the somatic motor prestin, our previous work has shown that salicylate directly affects the mechanics of the stereocilia bundle, increasing the amplitude of sound-evoked deflections, likely by reducing the bundle’s bending stiffness [16]. While this effect on the fast, cycle-by-cycle motion of the bundle is distinct from the slow (seconds-long) somatic contraction measured in the present study, a change in the mechanical load at the cell’s apical pole could indirectly influence the somatic response. Nevertheless, our central finding—that disabling the OHC motor with salicylate accelerates the slow somatic contraction while dramatically slowing the electrical potential—is most directly explained by the unmasking of a passive mechanical response in the cell body. The effect on stereocilia represents a parallel action of the drug rather than a direct cause of the accelerated somatic motility we report here.

The protective mechanism we describe should be distinguished from the fast adaptation of the mechanoelectrical transducer (MET) channel, which operates on a sub-millisecond timescale at low stimulus levels [17–19]. The process we have identified is different: it operates on a timescale of milliseconds to seconds, is engaged by intense sound, and is driven by the somatic motor protein prestin. Our results therefore provide a direct kinetic and molecular explanation for the foundational observations of slow, reversible contractions of the organ of Corti during acoustic overstimulation [8, 20]. The remarkable consistency between our measurements and historical reports validates our new technique. For instance, our measured peak axial OHC displacement (≈ −7.2 µm) aligns exceptionally well with foundational studies reporting organ of Corti width contractions of 9 µm [20] and Hensen’s cell displacements of 5 to 12 micrometers [8]. On the molecular level, we confirmed that this slow contraction is driven by a rise in intracellular Ca^2+^, as it was significantly attenuated by the chelator EGTA—functionally linking our results to the known effects of acoustic overstimulation [20]. Critically, whereas prior methods could only resolve the recovery from these contractions over tens of minutes, our AI-enhanced approach resolves their onset kinetics on a seconds timescale (*τ* = 3.0 s in control), revealing a dynamic interplay that was previously beyond temporal reach. This comparison demonstrates that our method accurately captures the same core phenomenon observed historically while adding a new, previously unmeasurable dimension of high-speed kinetic information.

Our work complements recent findings that identify a parallel mechanism for slow cochlear contractions mediated by the TRPA1 channel [21]. In that pathway, TRPA1 acts as a sensor for oxidative stress byproducts in non-sensory Hensen’s cells, initiating a long-lasting Ca^2+^ signal that propagates to and causes contractions in pillar and Deiters’ cells [21]. The prestin-dependent mechanism we describe is fundamentally different: it is intrinsic to the OHC itself, is triggered directly by membrane potential rather than by chemical damage signals, and operates on a faster timescale (seconds vs. minutes) to directly sharpen the cochlear electrical response. Together, these studies suggest the cochlea has evolved at least two distinct, slow-acting mechanical systems for protection: a TRPA1-mediated pathway in supporting cells that manages the response to cellular damage over longer periods, and a prestin-mediated system within the OHC that provides immediate kinetic control of the electrical response during overstimulation. Our findings provide a kinetic explanation for previously observed phenomena. The organ-level contractions described by Fridberger and Flock et al. [9, 22] can now be understood not just as a structural change, but as a process underpinned by this OHC kinetic control. While supporting cells contribute to the overall displacement (Supplementary Fig. S7), our results show that the kinetic sharpening of the receptor potential is specifically and exclusively tied to the OHC’s intrinsic, prestin-dependent motility. Furthermore, our results complement the recent discovery that the tectorial membrane’s (TM) Ca^2+^ reservoir is depleted during loud sound [10]. We show that the OHC kinetic control mechanism operates in parallel to this ionic regulation; indeed, when we chelated Ca^2+^ with EGTA, the OHC contraction was significantly attenuated, but the electrical potential persisted, albeit altered. This suggests that the cochlea employs at least two distinct, concurrent strategies to manage acoustic stress: ionic homeostasis mediated by the TM, and kinetic control mediated by the OHC motor.

This work also provides a mechanism for the recently established polarity switch of the summating potential (SP) from positive in a healthy state to negative under acoustic stress [11]. The inverse kinetic relationship we have uncovered governs the cochlea’s transition into this stressed, negative-SP state. The OHC’s contraction is what enables the system to maintain a finely-tuned operating point; without it, the system defaults to a sluggish electrical regime characteristic of injury. This entire process occurs within a tightly integrated mechanical system where even IHC stereocilia are physically coupled to the TM [7], reinforcing the need for mechanisms to precisely manage forces across the cochlear partition.

While our experiments were conducted in the cochlear apex, the fundamental mechanisms—prestin-dependent motility and the role of the SP—are conserved throughout the cochlea. Indeed, OHC somatic motility has been observed at frequencies of tens of kilohertz in more basal regions [23, 24]. Future studies using techniques like optical coherence tomography could explore how the exact time constants of this mechano-electrical interplay vary tonotopically. Furthermore, the discovery that salicylate can uncouple the normal kinetic relationship, allowing for a faster mechanical response at the expense of electrical fidelity, opens intriguing avenues for investigating therapeutic strategies for noise-induced hearing loss.

## Conclusion

In conclusion, we have demonstrated that the OHC’s slow somatic contraction is not a byproduct of intense stimulation but a central element of a sophisticated control system. This mechanism repurposes the molecular machinery of low-level amplification for a different function at high sound levels: protection and control. By acting as a mechanical damper on its own voltage, the OHC preserves the speed and temporal fidelity of the cochlear electrical response. This work reveals a dynamic and previously unknown layer of cellular control that is essential for hearing under demanding acoustic conditions. **Finally, given the established link between intense sound, negative SP polarity, and injury, the inverse kinetic relationship described here may help explain early changes in temporal fidelity during acoustic overexposure and suggests targets for protecting against noise-induced hearing loss.**

## Materials and Methods

### Animal Preparation and Experimental Setup

All procedures were conducted according to ethical guidelines approved by the Regional Ethics Committee in Linköping, Sweden (DNR 5111-2019, DNR 01122-2020 ID2529, DNR 19058-2021 ID4016). Guinea pigs (Cavia porcellus, 2–5 weeks old, either sex) were handled as previously described [7, 11, 25]. Briefly, after anesthesia and decapitation, the temporal bone was isolated and the bulla opened. This procedure yields an isolated temporal bone preparation, a configuration that has been previously illustrated [26]. The middle and inner ear were immersed in oxygenated minimum essential medium with Earle’s balanced salts (MEM, Gibco). Under a stereomicroscope, the cochlea was exposed and two small openings were made: one at the base for perfusion and one at the apex for imaging and recording. Middle ear immersion attenuated sound stimuli by approximately 25 dB SPL, and all reported intensities are corrected for this. The preparation was mounted in a custom rotatable holder on the stage of the microscope within a light-isolated Faraday cage on a vibration-dampening table.

### Electrophysiology and Pharmacology

Glass microelectrodes (1.5 mm outer diameter) were fabricated using a standard puller (P-1000, Sutter Instrument) and beveled at a 20*^◦^* angle to an impedance of approximately 3 MΩ. Electrodes were filled with artificial endolymph (1.3 mM NaCl, 31 mM KHCO_3_, 128.3 mM KCl, 0.023 mM CaCl_2_; pH 7.4; 300 mOs*/*kg) and mounted on a computer-controlled micromanipulator. **To ensure precise co-localization of the electrical and mechanical measurements, the preparation was maintained in a standardized cross-section orientation that is optimal for both imaging and electrode access.** Under continuous confocal imaging guidance, the electrode was inserted through Reissner’s membrane into the scala media of the cochlear apical turn, a region with a best frequency of approximately 180 Hz [11]. **Once the electrode was in place, a placement consistent with that shown previously [11], the confocal imaging plane (Z-plane) was optimized at this precise location and maintained throughout the experiment, guaranteeing that the recorded cellular mechanics corresponded directly to the source of the electrical potentials.**

For local pharmacological manipulation, 100 µM EGTA or 5 µM DHS was included in the recording electrode’s endolymph solution and delivered directly into the scala media via brief, low-pressure ejection (<4 psi), a method confirmed not to cause hydrostatic artifacts (Supplementary Fig. S11; Supplementary Movie 3). For salicylate treatments, 10 mM salicylate was added to the perfusion medium and allowed to equilibrate [16, 26, 27]. Cochlear microphonics were monitored to ensure preparation stability before and after drug administration.

### AI-Powered Droplet Volume Quantification

To validate that local drug delivery via picospritzer did not introduce significant hydrostatic pressure artifacts (Supplementary Fig. S11; Supplementary Movie 3), the ejected droplet volume was quantified using an *ex vivo* experiment. For this validation, a standard beveled glass microelectrode was positioned in air and imaged using a CMOS camera (CS165CU/M, Thorlabs). Droplets were ejected using the standard experimental parameters (4 psi for 1 second) while time-series images were recorded (1 frame/second). This procedure was repeated 11 times. The recorded image series were then processed using a custom AI segmentation pipeline. We implemented a U-Net architecture [28] using PyTorch [29] and the Segmentation Models PyTorch library [30]. The model employed a ResNet34 encoder with ImageNet pre-trained weights and included squeeze-and-excitation attention modules [31] in the decoder path to enhance feature representation. To improve model robustness, data augmentation was performed using the Albumentations library [32], including geometric transformations (e.g., flips, rotations) and photometric adjustments.

The model was trained on manually annotated droplet images for up to 70 epochs, with early stopping based on the validation Intersection over Union (IoU). Training utilized a combined loss function weighted between Dice loss (70%) and binary cross-entropy (BCE) loss (30%), the AdamW optimizer [33], and a cosine annealing learning rate schedule [34]. Mixed precision training [35] was employed to accelerate computation on CUDA-enabled GPUs. The model showed excellent performance and generalization, achieving a best validation IoU of 0.951 with a final train-validation loss gap of only 0.0127 (Supplementary Fig. S12).

For the final analysis, the maximum volume in each of the 11 recordings was identified (Supplementary Fig. S11e). The mean maximal volume across experiments was 37.2 ± 2.8 nL (mean ± SD). This volume represents less than 0.36% of the total guinea pig cochlear fluid volume (10.4 µL) and approximately 2.5% of the cochlear endolymph volume (1.5 µL) [36], confirming that the ejected volume is minimal and unlikely to cause significant hydrostatic artifacts. Given that endolymph constitutes the immediate dilution compartment, the locally applied pharmacological agents were rapidly diluted by a factor of at least 40, ensuring that effective concentrations remained localized and transient.

### Synchronized Data Acquisition

A custom LabVIEW (National Instruments) interface precisely synchronized acoustic stimulation, electrical recording, and confocal imaging (Fig. 1). Tone bursts at 160 Hz (1000 ms total duration) were generated. This frequency was chosen to be near the best frequency of the apical recording location (≈180 Hz) to ensure a strong and targeted cellular response, a strategy consistent with established protocols [10, 11]. The bursts consisted of a 720 ms active sound period flanked by 140 ms silent intervals. A pre-programmed sequence of 412 bursts stepped through different sound intensities (0 dB, 70 dB, 105 dB SPL).

Electrical signals were amplified (NPI BA-01X, 10× gain) and digitized at 10 kHz (NI USB-4431). High-speed imaging was performed on a Zeiss LSM 980 confocal microscope in reflection mode (488 nm laser) using a 40×, 0.8 NA water immersion lens, acquiring 512×512 pixel images in 630 ms. Each image acquisition was triggered to coincide with the central portion of the 720 ms active sound period, ensuring direct temporal alignment between mechanical and electrical data.

### Data Processing and Analysis

#### Electrical Signal Processing

To facilitate real-time monitoring of the experiment’s progress, the acquisition software stored electrical data in a hybrid format: 90% of the raw electrical waveform points were stored directly, while 10% were captured as a down-sampled rolling average. In post-processing, a continuous, full-resolution dataset was reconstructed from this format in MATLAB (MathWorks) by interpolating the rolling-average points using the interp1 function. Subsequent analysis of this reconstructed dataset was conducted in Python using the SciPy and NumPy libraries. For each recording, a linear baseline was calculated by applying a first-order polynomial fit (numpy.polyfit) to the pre-stimulus (0-140 ms) and post-stimulus (875-1000 ms) silent periods and then subtracted from the entire waveform. The direct current summating potential (SP) was isolated from this baseline-corrected signal by applying a moving average filter (scipy.ndimage.uniform_filter1d) with a window size corresponding to one cycle of the 160 Hz stimulus (≈6.3 ms, 63 samples at 10 kHz). The alternating current (AC) component was defined as the baseline-corrected, unfiltered waveform.

#### Image Processing and AI-Powered Segmentation

To maximize signal-to-noise, raw reflected-light confocal image series first underwent a two-stage denoising process, using sequential 2D-wavelet denoising for spatial noise reduction, followed by FastDVDnet video denoising for temporal noise reduction [14, 37]. Mechanical responses were then quantified from these denoised image series using a custom-built active learning pipeline detailed below.

##### Deep Learning-Based Segmentation and Active Learning

Our study employed an active learning approach to develop a robust segmentation model, a strategy inspired by the code-free pipeline described by Pettersen et al. (2022) [38]. The process was initiated by manually annotating an initial dataset in open-source software [38], which served as the foundation for training our deep learning model. We initially defined nine distinct classes: a general background class, tectorial membrane (TM), border cells (BC), outer hair cells (OHC), Hensen’s cells (HeC), reticular lamina (RL), pillar cells (PC), inner hair cells (IHC), and Reissner’s membrane (RM). For segmentation clarity, the PC label depicts the outer pillar cell heads, while the IHC label represents the IHC/inner pillar complex. However, Reissner’s membrane (RM) was excluded from the final analysis due to its dynamic behavior, which caused it to transiently move out of the acquisition frame during intense sound stimulation, leaving eight classes in the final model. The core of our auto-annotation pipeline was a DeepLabV3Plus architecture with a ResNet18 encoder. To enhance model generalization, a comprehensive suite of data augmentation techniques was applied, including resizing, horizontal and vertical flipping, rotation, brightness and contrast adjustments, gamma correction, blurring, and grid distortion. The training process itself was iterative: the model was trained on the annotated set, used to predict segmentations on new, unannotated images, and these new auto-annotations were then manually corrected and added to the training set. This cycle of training, prediction, and correction allowed for the continuous expansion and refinement of our dataset while maintaining high-quality standards. Post-inference, results were refined using morphological operations (e.g., expansion and erosion) and an adaptive thresholding mechanism to identify high-quality annotations based on an Intersection over Union (IoU) score, typically ranging from 0.97 to 0.985.

##### Quality Control and Model Evaluation

A rigorous quality control protocol was essential to the pipeline. Each auto-annotation was automatically checked to ensure all required anatomical classes were present with plausible sizes and spatial relationships (e.g., ensuring the OHC class was not positioned above the RL). To further refine the dataset, we analyzed metrics for each segmented object, including centroid coordinates, length, width, and area. A three-frame rolling window analysis helped identify temporal outliers or inconsistencies, flagging them for manual re-annotation. This systematic quality control ensured the integrity and accuracy of the final training dataset.

To evaluate the final model, the fully annotated dataset was split into training (85%) and validation (15%) sets. Crucially, this split was performed at the time-series level, ensuring that no frames from a given time-series appeared in both sets, thus preventing data leakage and providing a robust, independent evaluation. The final model achieved high performance, with a mean IoU of 0.913 and a mean pixel accuracy of 0.987 on unseen test data, demonstrating its ability to generalize and accurately delineate cochlear structures. The model’s performance is further detailed in the confusion matrices (Supplementary Fig. S14). The complete AI-powered segmentation pipeline is visualized in Supplementary Fig. S13.

##### Metric Quantification

Following segmentation, morphological metrics—including X and Y centroid coordinates, area, lengths, and widths—were computed for each structure in every frame. The precision of these spatial measurements is grounded in the high pixel resolution of the LSM 980 microscope and our robust AI-driven quantification. In a recent touch receptor stimulation study, our lab used AI-based segmentation and pixel-based motion analyses alongside optical coherence tomography (OCT)—a gold standard for measuring cochlear mechanical responses—to provide a high-fidelity characterization of stimulus mechanics and found a strong agreement between these techniques with OCT [39]. For the 412-image series, a baseline correction was performed using the median of the initial 63 data points recorded at 0 dB SPL. For the high-speed, time-resolved series (Fig. 5), data were first baseline-corrected using their median value, followed by temporal smoothing with a 3-point moving average to reduce noise while preserving underlying dynamics.

### Kinetic and Statistical Analysis

All kinetic and statistical analyses were performed in Python using the NumPy, SciPy, and Matplotlib libraries. The kinetics of electrical (SP) and mechanical (OHC area change) responses were characterized by fitting first-order exponential models of the form *y*(*t*) = *c* + *a* · *e^−t/τ^* to the mean traces using the Levenberg-Marquardt algorithm implemented in `scipy.optimize.curve_fit`. For fitting stability, time vectors were often normalized to a [0, 1] range, and the resulting rate constants were scaled back to obtain the time constant (*τ*) in physical units (ms or s). Fits were considered reliable only if the coefficient of determination *R*^2^ ≥ 0.8.

Data are presented as mean ± s.e.m. For all cochlear experiments, n refers to the number of independent cochlear preparations, which corresponds to the number of animals used. For group comparisons, normality of the data distributions was first assessed using the Shapiro-Wilk test (`scipy.stats.shapiro`). For normally distributed data, statistical significance (*p <* 0.05) between two groups was determined using an independent samples t-test (`scipy.stats.ttest_ind`), and among multiple groups using a one-way analysis of variance (ANOVA) (`scipy.stats.f_oneway`). For non-normally distributed data, we used the non-parametric Mann-Whitney U test for two-group comparisons and the Kruskal-Wallis test for multiple-group comparisons, implemented with the `scipy.stats.mannwhitneyu` and `scipy.stats.kruskal` functions, respectively. The temporal relationship between normalized electrical and mechanical signals was quantified using the Pearson correlation coefficient (`scipy.stats.pearsonr`).

## Author Contributions

A.F. developed the automated data acquisition, aligning it with the established acquisition paradigm and contributed with expertise in cochlear mechanics. P.H. conceptualized and designed the study, as well as conducted the experimental procedures. The segmentation of the hearing organ into structures of interest was a joint effort between P.H. and H.S.P., with H.S.P doing more work. The analysis of the neural networks’ output data was a joint effort by H.S.P. and P.H., working from datafiles generated by H.S.P. P.H. took charge of analyzing the electrical recordings and performing statistical quantifications; PH also performed the U-Net/ResNet34 segmentation of the injection droplet experiments and the related volume analysis. The manuscript was written by P.H., with input from the co-authors.

## Competing Interest Statement

The authors declare no competing interests.

## Acknowledgments

P.H. acknowledges the grants FB23-0003 FB24-0005 from the Swedish Tysta skolan foundation and M3 from the Torsten Söderberg foundation. H.S.P. acknowledges multiple grants from the Research Fund for the Center for Laboratory Medicine, St. Olavs hospital.

## Data Availability Statement

All data supporting the findings of this study are available within the manuscript and its Supplementary Information files. Raw datasets are available from the corresponding author upon reasonable request.

## Supplementary Figures

**Supplementary Figure S1:**
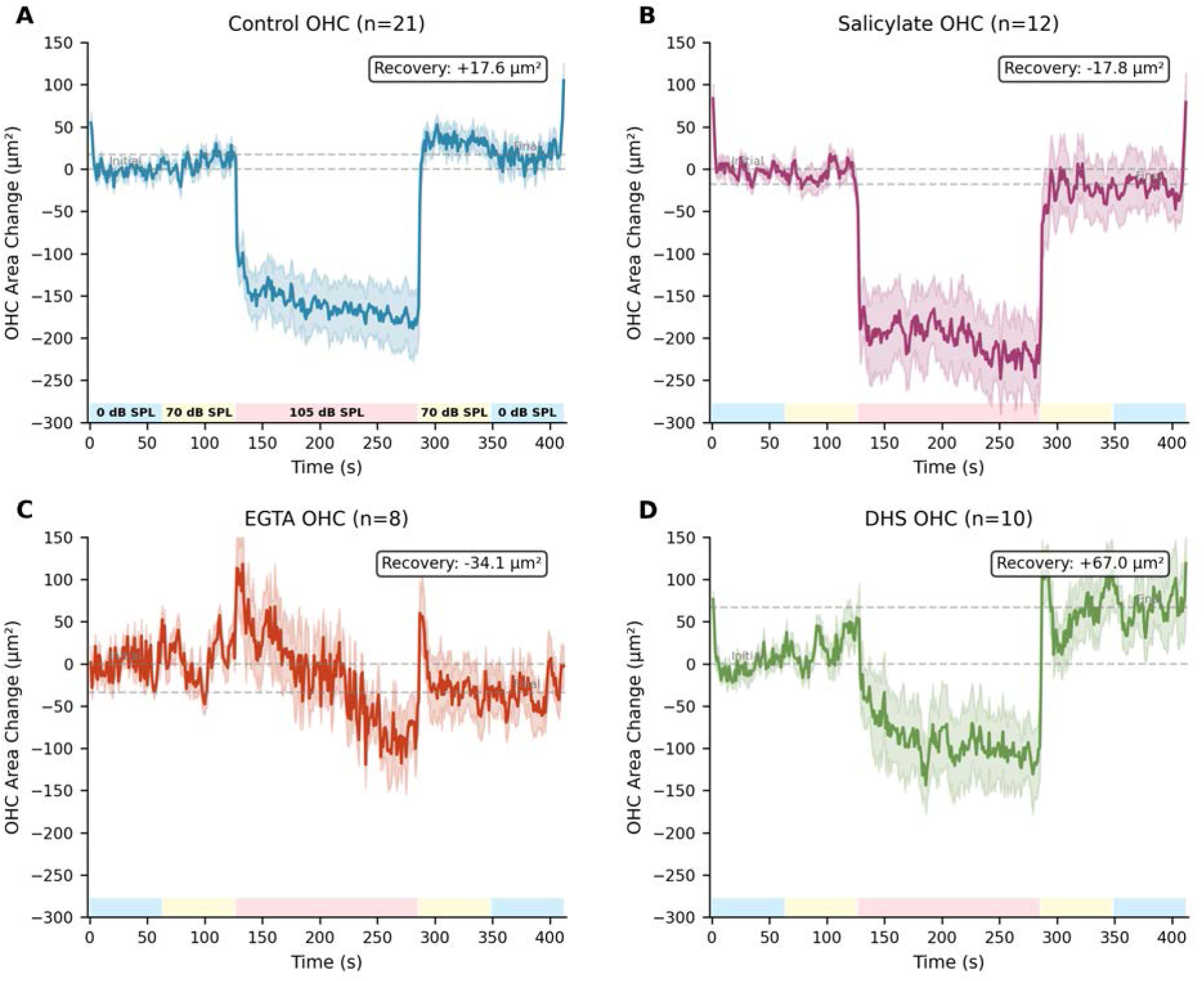
Complete OHC response time course with recovery analysis. Complete time series showing OHC area changes for (**a**) Control (*n* = 21), (**b**) Salicylate (*n* = 12), (**c**) EGTA (*n* = 8), and (**d**) DHS (*n* = 10) conditions. After exposure to 105 dB SPL, control cells show a net swelling during the recovery period (+17.6 *µm*^2^), whereas salicylate and EGTA-treated cells exhibit residual shrinkage (−17.8 *µm*^2^ and −34.1 *µm*^2^, respectively). DHS-treated cells show a more pronounced swelling (+67.0 *µm*^2^).

**Supplementary Figure S2:**
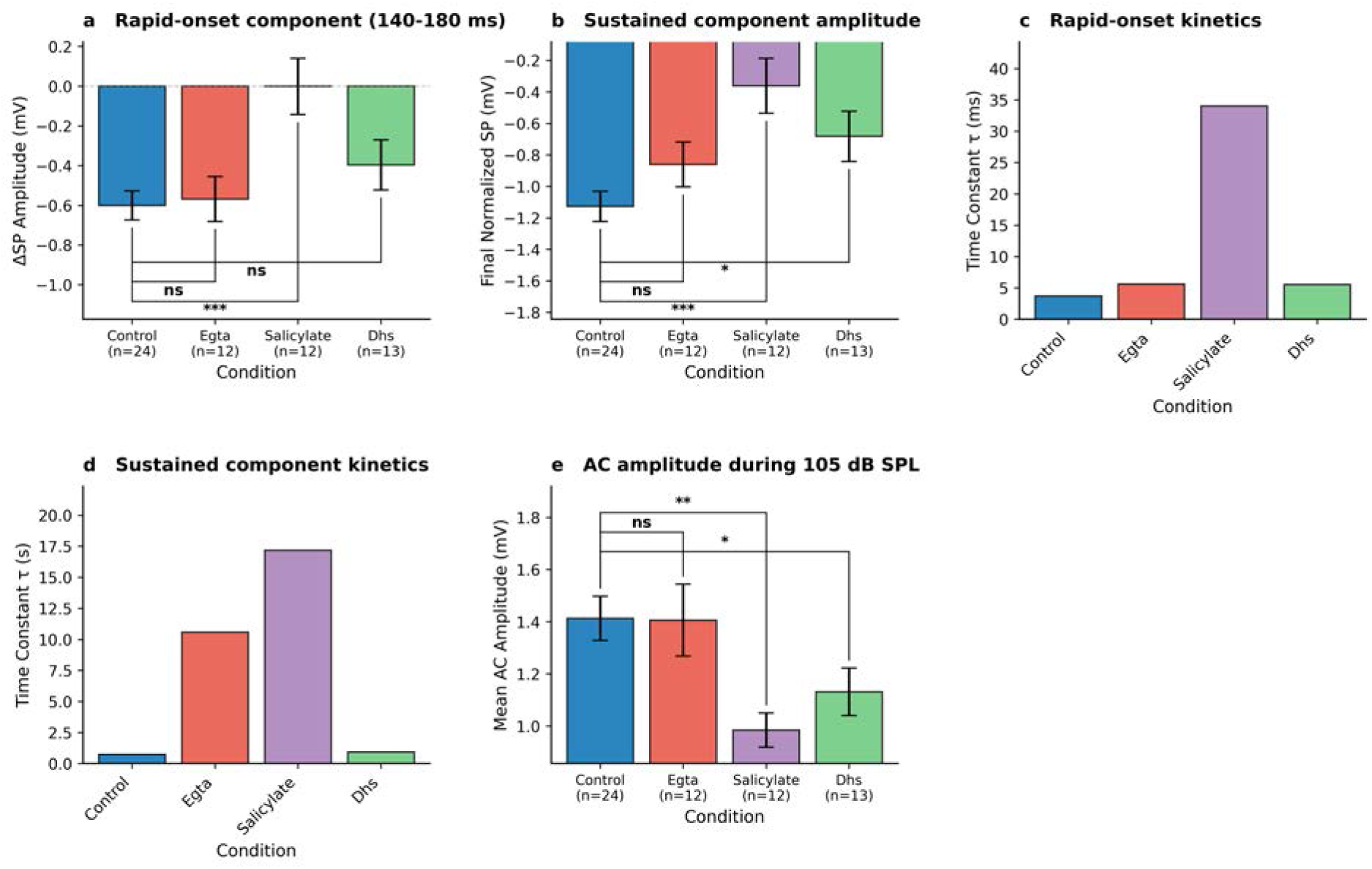
Summary of pharmacological effects on cochlear electrical responses. **a,** The rapid-onset component, measured as the change in SP amplitude from 140-180 ms during stimulation. Salicylate significantly attenuated this component compared to control (*p* = 0.0002). **b,** The sustained component amplitude, measured as the final normalized SP value. Both salicylate (*p* = 0.0002) and DHS (*p* = 0.0156) significantly reduced the amplitude relative to control. **c,** Time constants (*τ*) for the rapid-onset component. The kinetics were dramatically slowed by salicylate (*τ* = 34.0 ms) compared to control (*τ* = 3.7 ms). **d,** Time constants for the sustained component. Both EGTA (*τ* = 10.6 s) and salicylate (*τ* = 17.2 s) substantially slowed the decay kinetics compared to control (*τ* = 0.7 s). **e,** Mean AC amplitude during the 105 dB SPL period. Amplitude was significantly reduced by salicylate (*p* = 0.0022) and DHS (*p* = 0.0419). All data are presented as mean ± s.e.m. Sample sizes (n) are indicated in panel (a). Statistical significance shown is from pairwise comparisons to the control group: ns = not significant, ∗*p <* 0.05, ∗ ∗ *p <* 0.01, ∗ ∗ ∗*p <* 0.001.

**Supplementary Figure S3:**
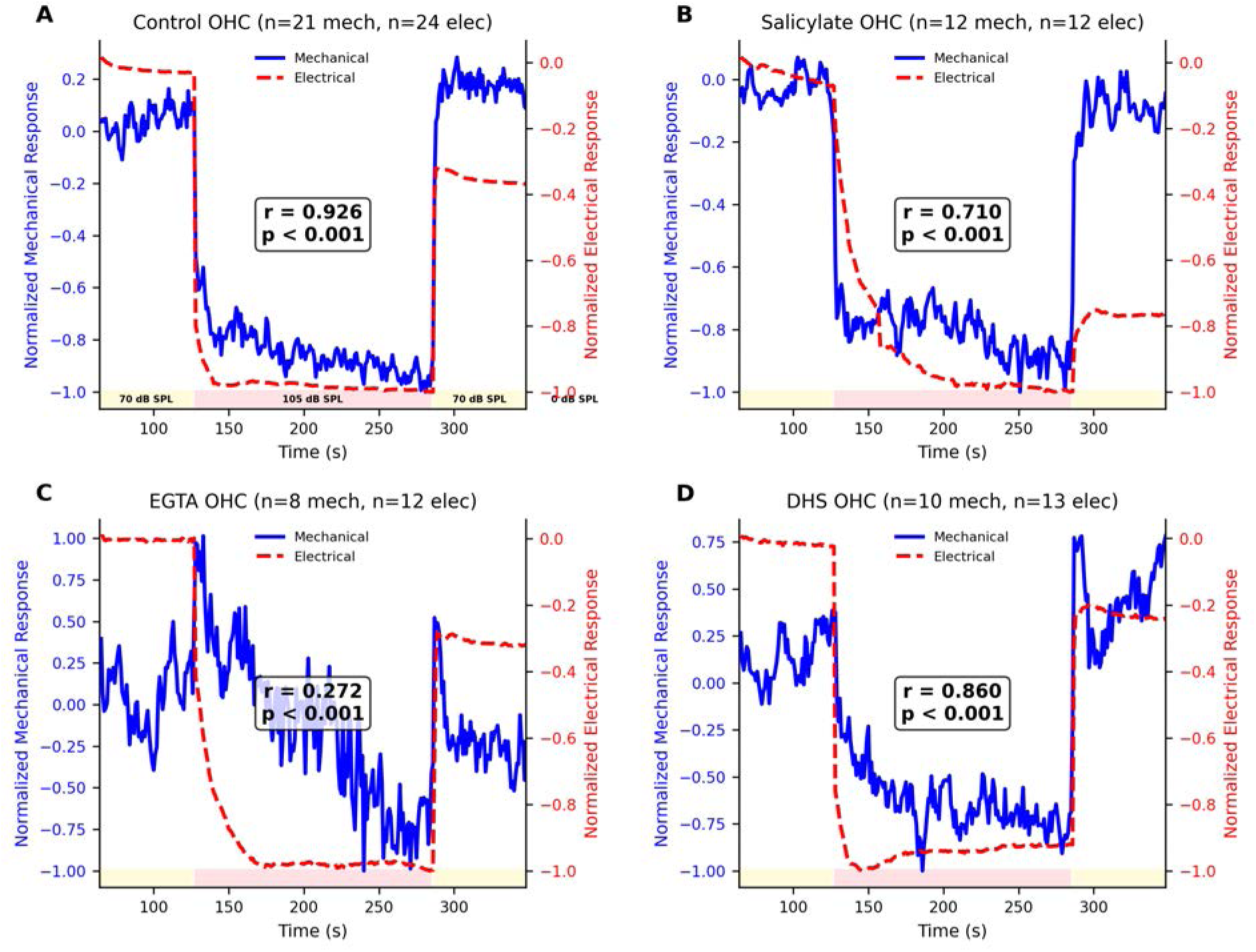
Electrical-mechanical coupling remains strong despite altered kinetics. Normalized electrical (red dashed lines) and mechanical (blue solid lines) responses are overlaid for (**a**) Control, (**b**) Salicylate, (**c**) EGTA, and (**d**) DHS conditions. Pearson correlation coefficients (*r*) confirm a strong temporal correlation in the control (*r* = 0.926) and salicylate (*r* = 0.710) conditions, indicating that the mechanical contraction remains tightly coupled to the stimulus-evoked potential even when its kinetics are altered.

**Supplementary Figure S4:**
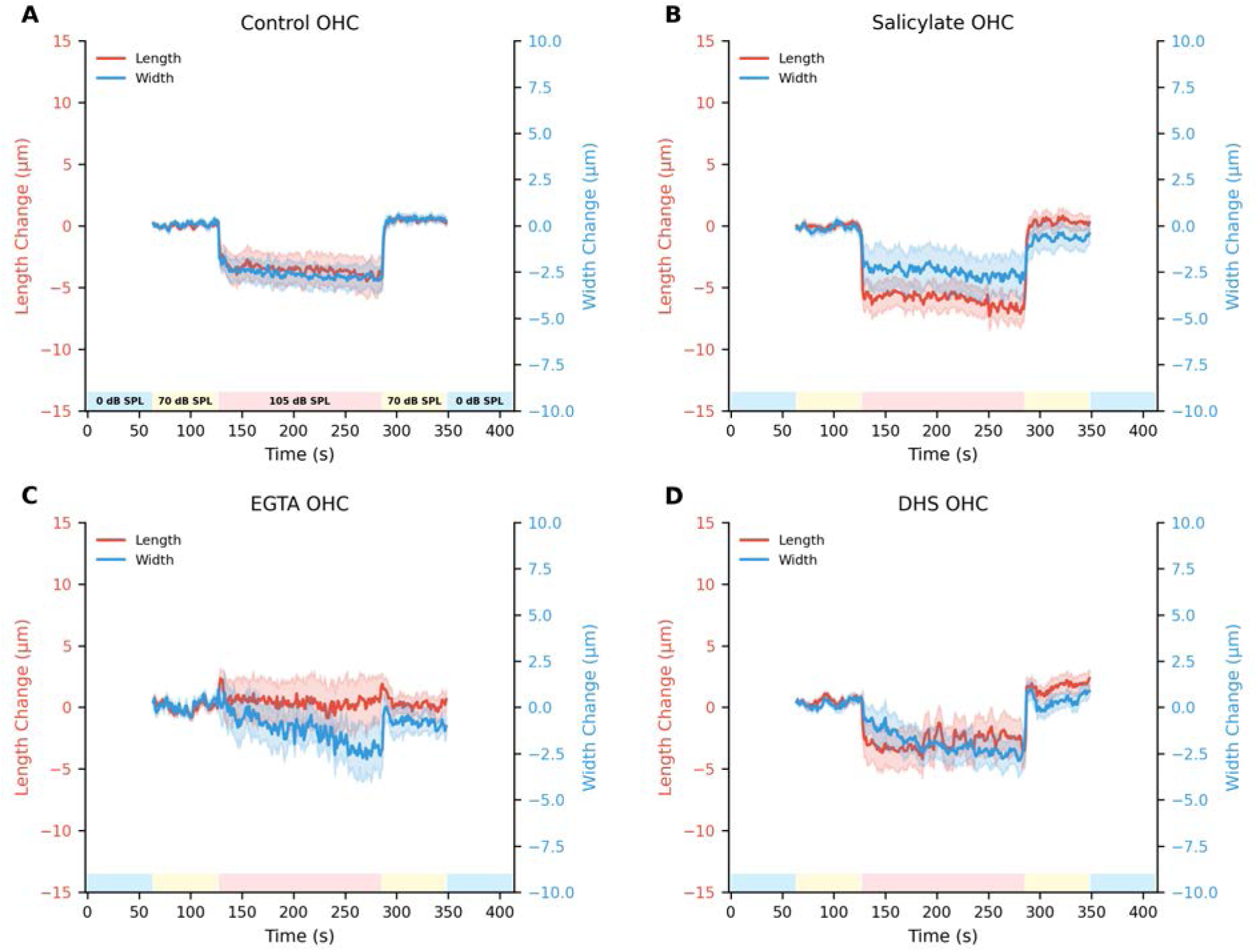
OHC morphological changes are anisotropic. Simultaneous measurement of OHC length (red, left axis) and width (blue, right axis) for (**a**) Control, (**b**) Salicylate, (**c**) EGTA, and (**d**) DHS conditions. The differential changes reveal the anisotropic nature of OHC contraction, which is dominated by a decrease in cell length along its major axis.

**Supplementary Figure S5:**
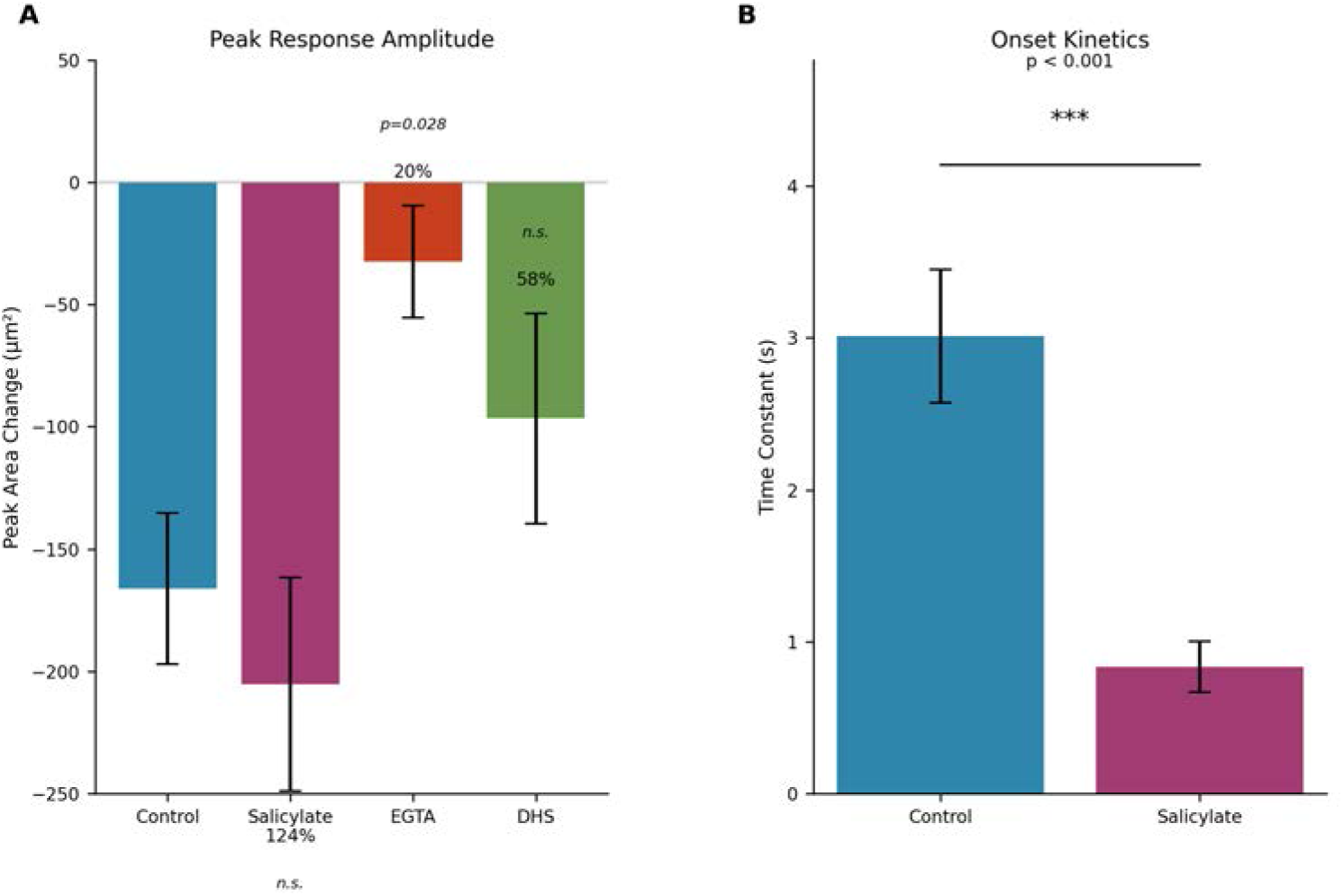
Summary of pharmacological effects on OHC mechanics. (**a**) Bar chart of peak OHC area change during 105 dB SPL stimulation. Control cells showed a mean contraction of 166 µm^2^, while salicylate-treated cells showed 205 µm^2^ (not statistically significant, *p* = 0.082). EGTA significantly reduced the contraction by 80% (*p <* 0.001), while DHS reduced it by 42% (*p <* 0.001). (**b**) Bar chart comparing onset time constants, showing the 3.6-fold acceleration of contraction kinetics when prestin is blocked with salicylate (*τ* = 0.84 s vs. 3.0 s, *p <* 0.001).

**Supplementary Figure S6:**
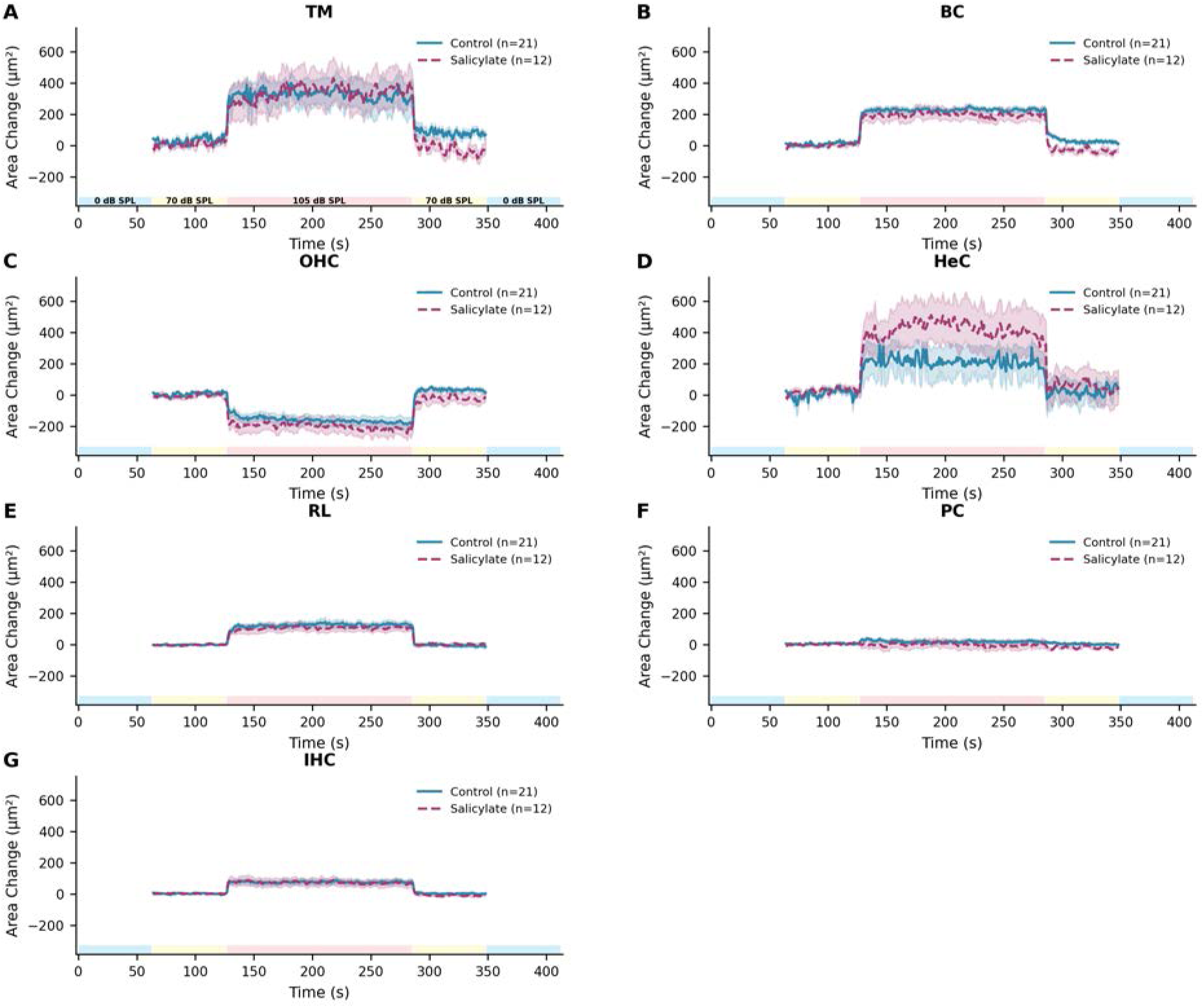
OHC-specific mechanical responses to sound stimulation. Comparison of area changes across seven different cochlear structures under Control (solid lines) and Salicylate (dashed lines) conditions. (**a**) Tectorial membrane, (**b**) Border cells, (**c**) Outer hair cells, (**d**) Hensen cells, (**e**) Reticular lamina, (**f**) Pillar cells, and (**g**) Inner hair cells. The large, sustained contraction during 105 dB SPL stimulation is unique to the OHCs.

**Supplementary Figure S7:**
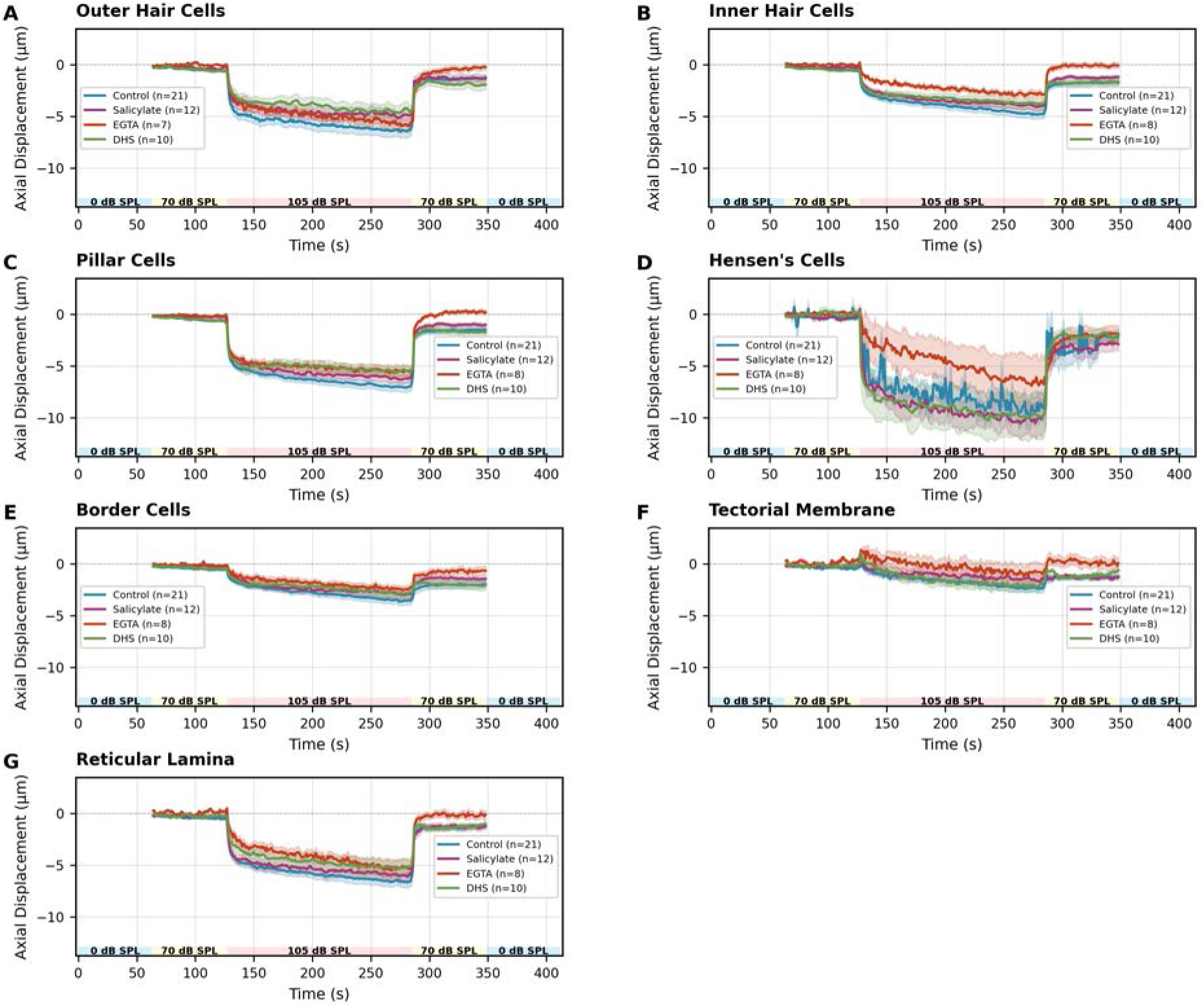
Coordinated axial displacement across cochlear cell types. Time course of axial displacement for seven cochlear structures under four pharmacological conditions. (**a**) Outer Hair Cells (OHCs) show a large displacement, reaching a peak of approximately −7.2 µm in control. (**b-g**) Inner hair cells (IHCs), pillar cells (PCs), and other supporting structures move in concert with the OHCs. Contrary to somatic contraction, the magnitude of this bulk displacement was not significantly altered by salicylate or DHS for any cell type, and only by EGTA for IHCs (see Supplementary Fig. S8).

**Supplementary Figure S8:**
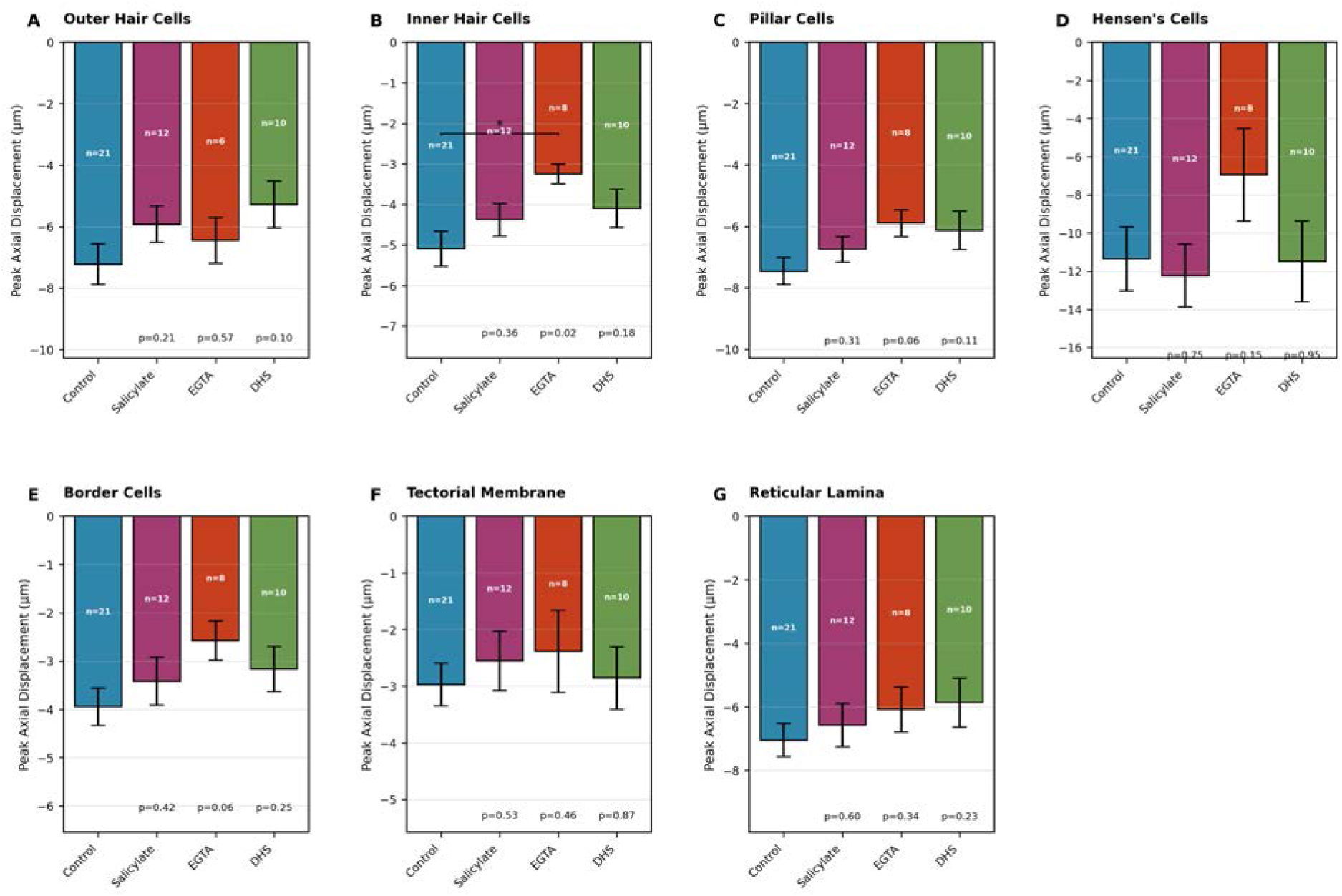
Statistical summary of peak axial displacement. Bar charts show peak displacement for each cell type across the four conditions. (**a**) For OHCs, no significant differences were found between control and drug treatments (ANOVA, *p* = 0.26). (**b**) For IHCs, displacement was significantly reduced only by EGTA treatment (t-test, *p* = 0.018). (**c-g**) No other statistically significant changes were observed for any other cell type.

**Supplementary Figure S9:**
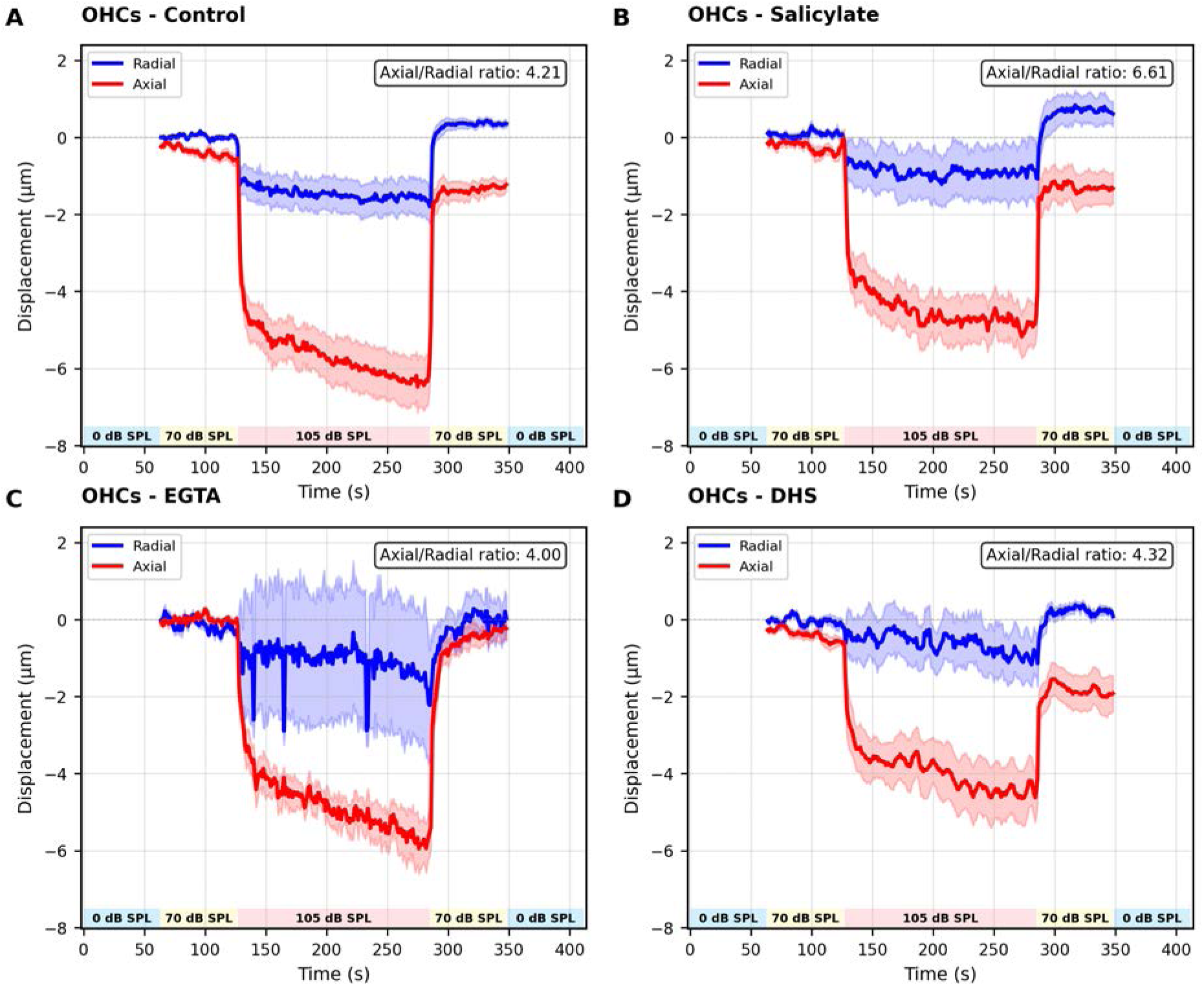
OHC displacement is predominantly axial. Simultaneous measurement of OHC radial (blue) and axial (red) displacement components for (**a**) Control, (**b**) Salicylate, (**c**) EGTA, and (**d**) DHS conditions. Axial displacement was consistently 4-to-7-fold larger than radial displacement, confirming that the dominant motion is in the axial direction, consistent with the OHC’s role in cochlear mechanics.

**Supplementary Figure S10:**
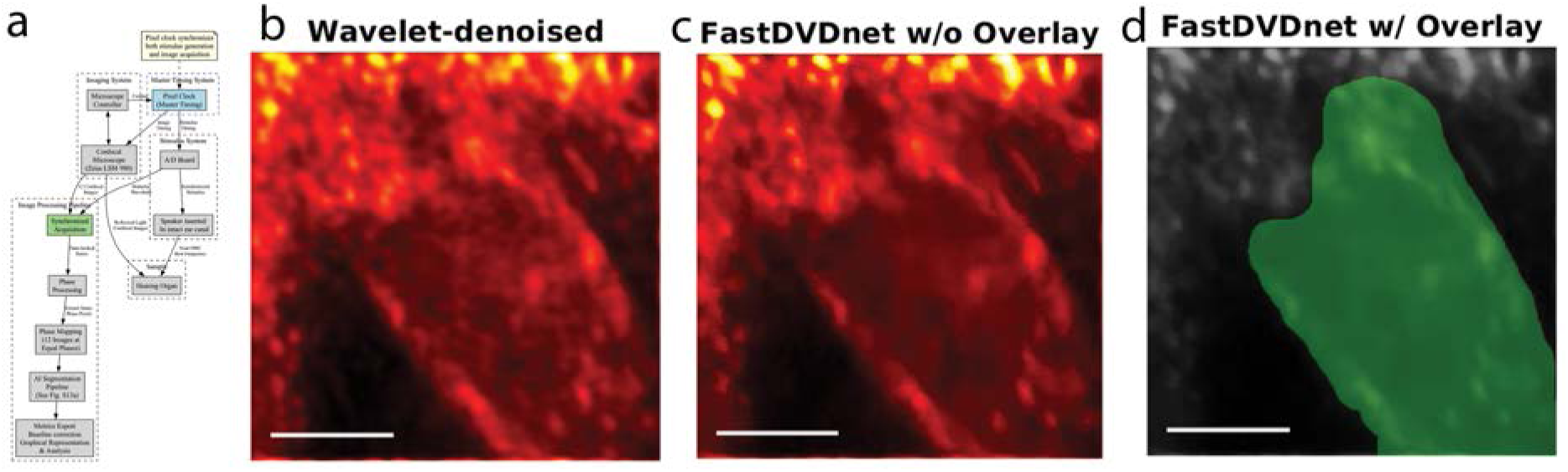
Methodology for high-speed imaging and analysis of cycle-by-cycle OHC mechanics. (**a**) Flowchart of the pixel-clock-synchronized imaging and analysis pipeline. (**b-d**) Demonstration of the image processing workflow. Raw images first undergo two-stage denoising using a 2D-wavelet filter, followed by FastDVDnet video denoising (image after denoising shown in **c**). Final segmentation is then performed using our general segmentation pipeline (Supplementary Fig. S13a), as shown by the mask overlay in (**d**). This specific cycle-by-cycle analysis employed a dedicated binary model (OHC vs. background), whose accuracy is validated in the confusion matrix in Supplementary Fig. S14a. In contrast, the analysis of the main experimental protocol used a more complex multi-class model, validated in Supplementary Fig. S14b. Scale bars are 5 µm.

**Supplementary Figure S11:**
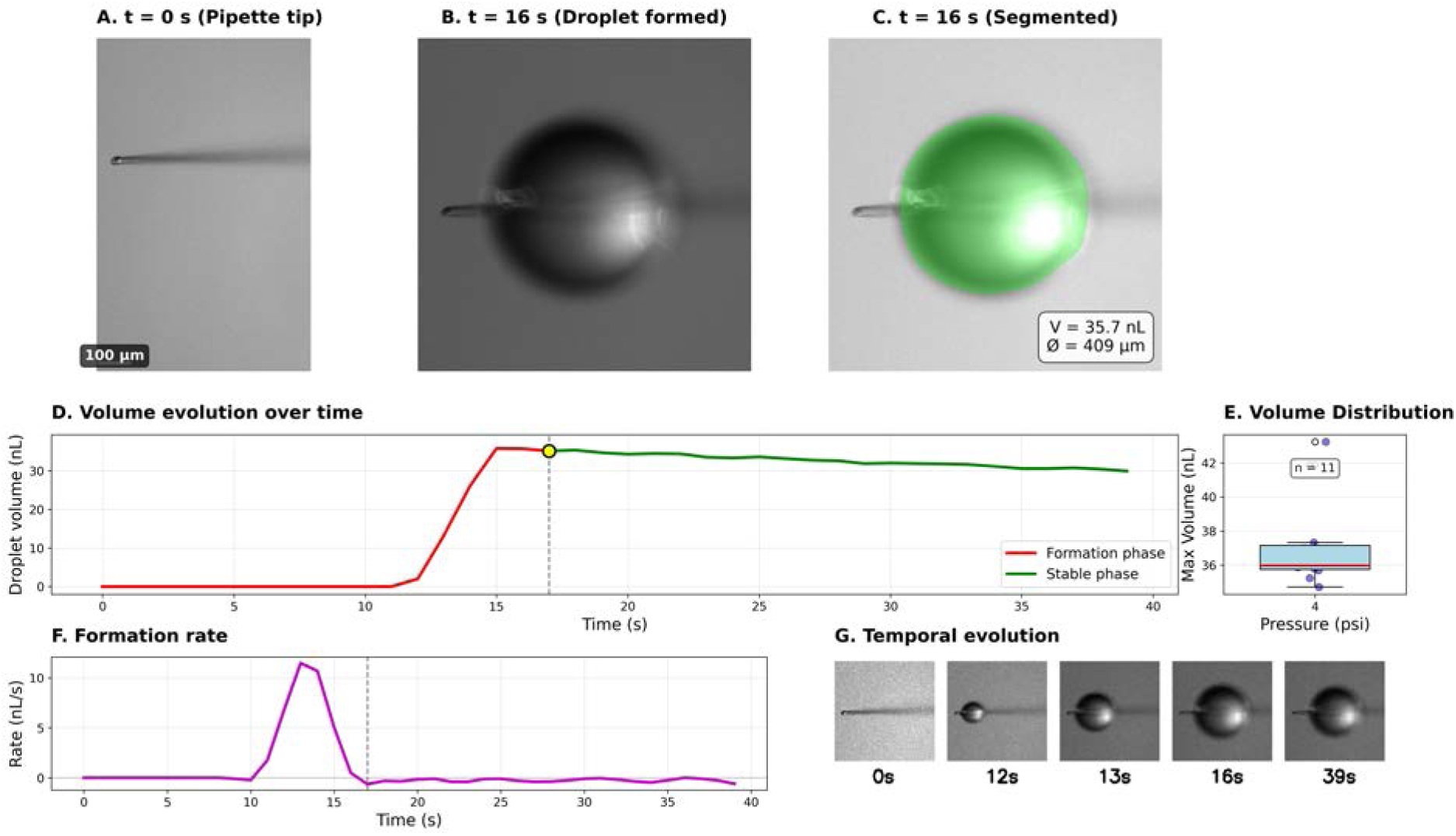
Validation of local drug delivery volume using AI-powered analysis. To confirm that local drug application via picospritzer does not introduce significant hydrostatic pressure, the ejected droplet volume was quantified in an *ex vivo* setup. (**a**) A pipette tip before injection (t = 0 s). (**b**) The formed droplet at t = 16 s. (**c**) The AI-segmented droplet with measured volume (V = 35.7 nL) and diameter (Ø = 409 µm). (**d**) Temporal dynamics of a representative droplet, showing the rapid formation phase (red) and a stable phase (green), with a maximum volume of 35.8 nL reached at 17 s. (**e**) Volume distribution from n = 11 independent experiments, showing high reproducibility. (**f**) The rate of volume formation, which peaked at 11.5 nL/s. (**g**) A time-lapse montage of key timepoints during droplet formation. Scale bar: 100 µm.

**Supplementary Figure S12:**
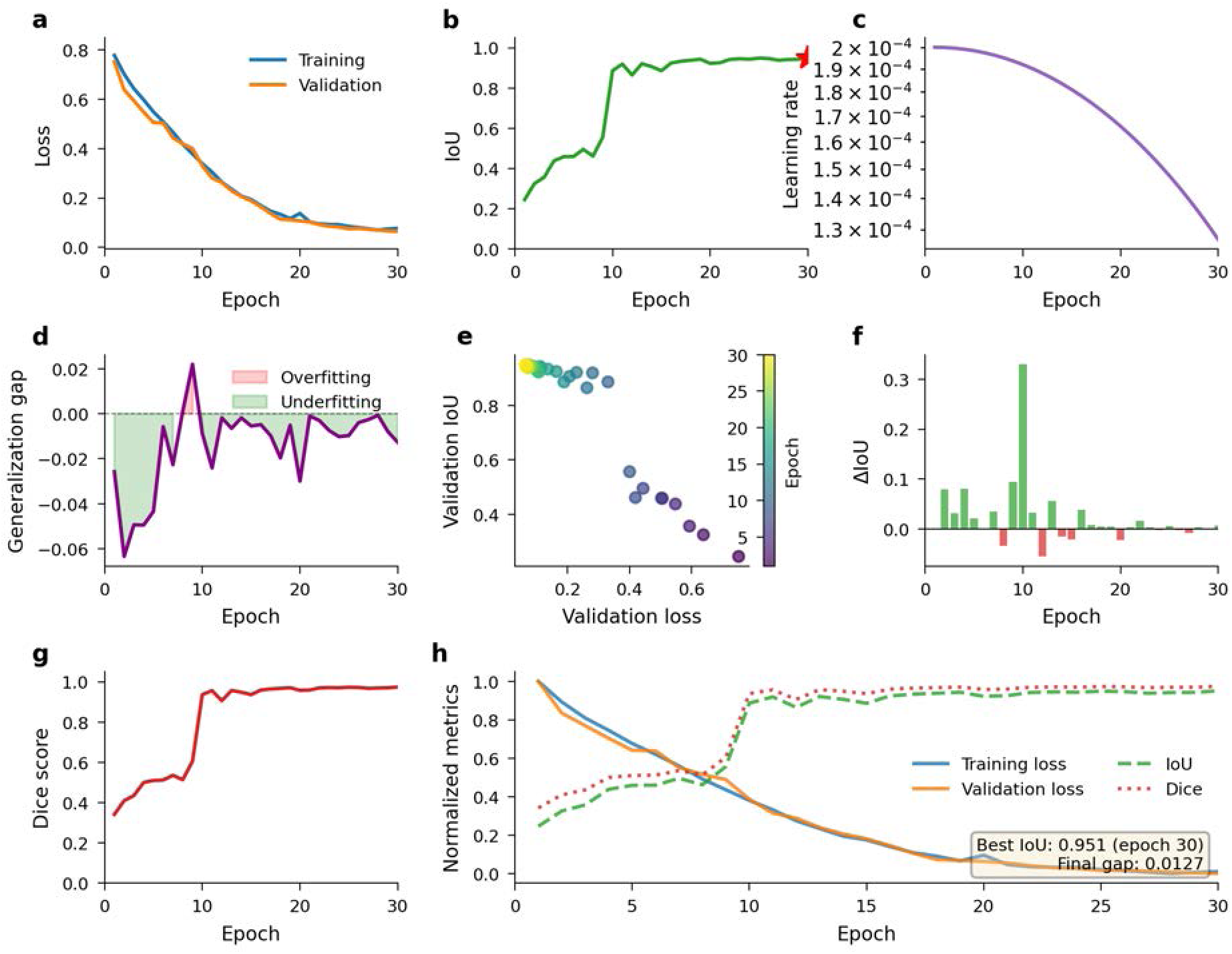
U-Net training dynamics for droplet segmentation. The figure summarizes the performance of the U-Net model over 30 training epochs. **a**, Training and validation loss curves show rapid convergence with the validation loss (orange) closely tracking the training loss (blue), indicating good generalization. **b**, Validation Intersection over Union (IoU), the primary performance metric, improves steadily and plateaus above 0.9, with the best performance of 0.951 achieved at epoch 30 (red star). **c**, The learning rate followed a cosine annealing schedule. **d**, The generalization gap (validation loss - training loss) remains close to zero, confirming minimal overfitting. **e**, A strong inverse correlation is observed between validation loss and IoU. **f**, Per-epoch IoU improvement was greatest in the early epochs. **g**, The Dice score, another segmentation metric, closely parallels the IoU, reaching a final value of 0.974. **h**, Normalized metrics confirm the synchronized convergence of all performance measures, with a final train-validation loss gap of only 0.0127, indicating excellent model generalization.

**Supplementary Figure S13:**
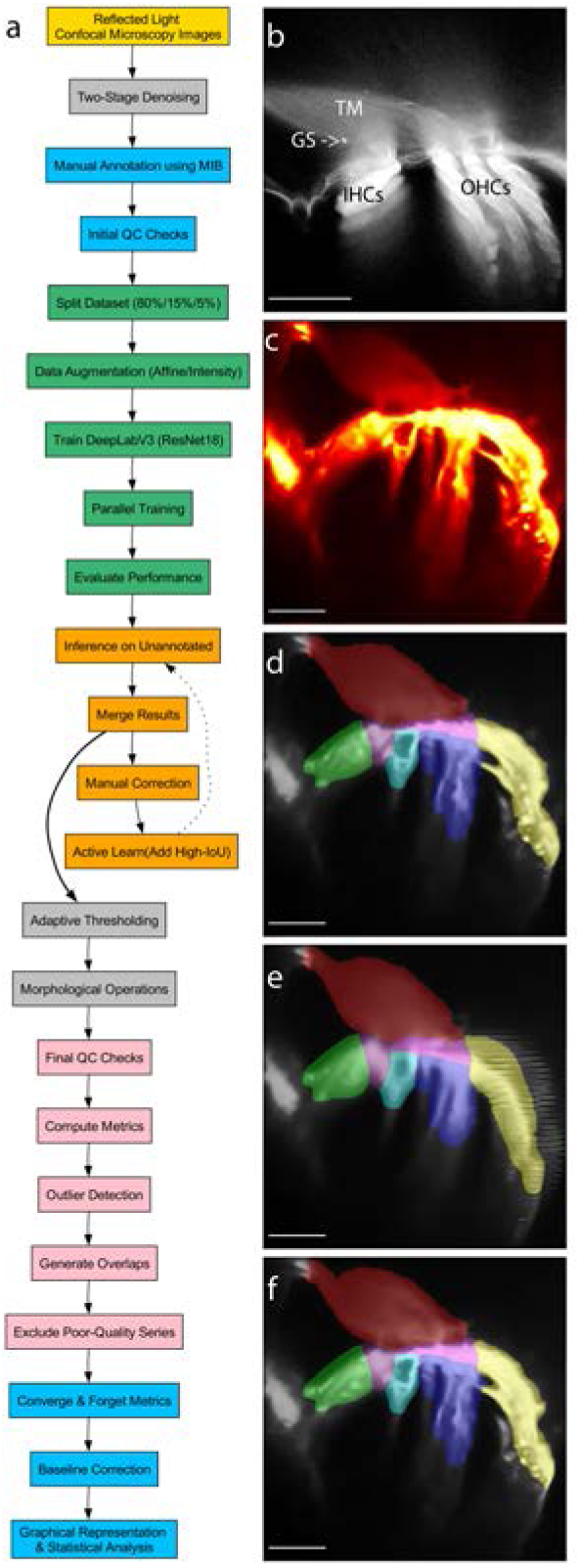
AI-powered segmentation pipeline for quantifying cochlear mechanics. **(a)** Flowchart of the custom active-learning pipeline. The process begins with two-stage denoising of raw confocal microscopy images, followed by manual annotation to train a DeepLabV3-based model, which is then iteratively improved through an active learning loop. **(b, c)** Representative images of the organ of Corti. For anatomical reference, a fluorescence image is shown in **(b)**. This panel shows key features including the tectorial membrane (TM), outer hair cells (OHCs), inner hair cells (IHCs), and the granular structures (GS) [7]. The high-speed mechanical analysis was performed on reflected-light images, an example of which is shown in **(c)**. The fluorescence imaging modality **(b)**, requiring over a minute per frame, is too slow to resolve the fast mechanical events captured with reflected-light imaging **(c)**, which was acquired in 630 ms. **(d-f)** Examples of the final segmented output, showing the precise tracking of cochlear structures before **(d)**, during **(e)**, and after **(f)** intense sound stimulation. The color code for the segmented structures, also visualized in the supplementary segmentation video, is as follows: tectorial membrane (TM, brown), border cells (BC, green), outer hair cells (OHCs, blue), Hensen’s cells (HeC, yellow), and reticular lamina (RL, clear magenta). For segmentation clarity, note that the **IHC** label (dark magenta) represents the **IHCs/inner pillar complex**, and the **PC** label (cyan) depicts the **outer pillar cell heads**. Scale bars, 50 µm.

**Supplementary Figure S14:**
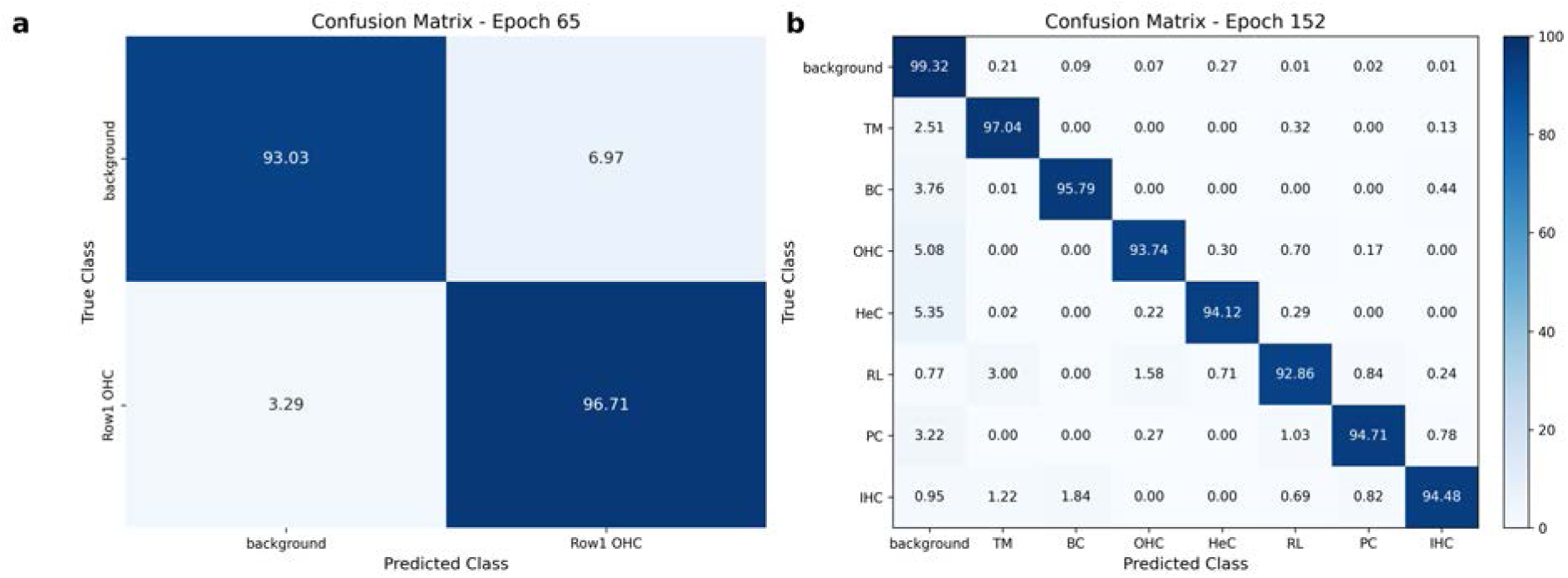
Performance of the DeepLabV3-ResNet18 segmentation model. (**a**) Normalized confusion matrix for the binary segmentation model (OHC vs. background) used for the high-speed, cycle-by-cycle mechanical analysis presented in Fig. 5 and Supplementary Fig. S10. The model achieves high accuracy, correctly classifying the OHC class over 96% of the time. (**b**) Normalized confusion matrix for the multi-class segmentation model used for the main mechanical analysis (e.g., Figs. 3-4 and associated supplementary figures). This model demonstrates high accuracy across eight distinct cochlear structures, with all diagonal values exceeding 92%. The low off-diagonal values in both matrices indicate minimal confusion between classes. The matrices compare predicted classes with ground-truth classes on a held-out test set not used during training. Abbreviations: TM, tectorial membrane; BC, border cells; OHC, outer hair cells; HeC, Hensen’s cells; RL, reticular lamina; PC, pillar cells; IHC, inner hair cells.

## Legends For Supplementary Videos

Supplementary Movie 1: Time-series acquired by reflected light confocal imaging (see Methods) after 2D-wavelet and FastDVDnet denoising. The full sequence shows the response to baseline, moderate (70 dB SPL), and intense (105 dB SPL) sound stimulation. (Filename: SupplementalMovie1.mp4)

Supplementary Movie 2: AI-powered segmentation of the cochlear structures shown in Supplementary Movie 1. The segmentation highlights the tectorial membrane, border cells, IHCs, pillar cells, reticular lamina, OHCs, and Hensen’s cells. (Filename: Supplemental_Movie2_Segmentation_Visualization.mov)

Supplementary Movie 3: Real-time quantification of ejected droplet volume using AI-powered segmentation. The video shows a side-by-side comparison of the original microscopy recording and the AI-segmented output, with real-time measurements. This movie corresponds to the analysis presented in Supplementary Fig. S11. (Filename: 4V4psi_final_publication_HQ.mp4)

## Notes

### Competing Interest Statement

The authors have declared no competing interest.

### Summary of Updates

Certain supplementary figure captions were incomplete.

